# Deciphering compromised speech-in-noise intelligibility in older listeners: the role of cochlear synaptopathy

**DOI:** 10.1101/2020.06.09.142950

**Authors:** Markus Garrett, Viacheslav Vasilkov, Manfred Mauermann, Pauline Devolder, John L. Wilson, Leslie Gonzales, Kenneth S. Henry, Sarah Verhulst

## Abstract

Speech intelligibility declines with age and sensorineural hearing damage (SNHL). However, it remains unclear whether cochlear synaptopathy (CS), a recently discovered form of SNHL, significantly contributes to this issue. CS refers to damaged auditory-nerve synapses that innervate the inner hair cells and there is currently no go-to diagnostic test available. Furthermore, age-related hearing damage can comprise various aspects (e.g., hair cell damage, CS) that each can play a role in impaired sound perception. To explore the link between cochlear damage and speech intelligibility deficits, this study examines the role of CS for word recognition among older listeners. We first validated an envelope-following response (EFR) marker for CS using a Budgerigar model. We then applied this marker in human experiments, while restricting the speech material’s frequency content to ensure that both the EFR and the behavioral tasks engaged similar cochlear frequency regions. Following this approach, we identified the relative contribution of hearing sensitivity and CS to speech intelligibility in two age-matched (65-year-old) groups with clinically normal (n=15, 8 females) or impaired audiograms (n=13, 8 females). Compared to a young normal-hearing control group (n = 13, 7 females), the older groups demonstrated lower EFR responses and impaired speech reception thresholds. We conclude that age-related CS reduces supra-threshold temporal envelope coding with subsequent speech coding deficits in noise that cannot be explained based on hearing sensitivity alone.

**Significance Statement:** Temporal bone histology reveals that cochlear synaptopathy (CS), characterized by damage to inner hair cell auditory nerve fiber synapses, precedes sensory cell damage and hearing sensitivity decline. Despite this, clinical practice primarily evaluates hearing status based on audiometric thresholds, potentially overlooking a prevalent aspect of sensorineural hearing damage due to aging, noise exposure, or ototoxic drugs—all of which can lead to CS. To address this gap, we employ a novel and sensitive EEG-based marker of CS to investigate its relationship with speech intelligibility. This study addresses a crucial unresolved issue in hearing science: whether CS significantly contributes to degraded speech intelligibility as individuals age. Our study-outcomes are pivotal for identifying the appropriate target for treatments aimed at improving impaired speech perception.

## Introduction

The cochlea, deeply embedded within the temporal bone, poses challenges for direct histological assessments of sensorineural hearing loss (SNHL) in living humans. While outer hair cell (OHC) deficits are typically diagnosed through indirect methods like pure-tone audiograms or distortion-product otoacoustic emissions (DPOAEs), damage to auditory nerve fiber (ANF) synapses between inner hair cells (IHCs) and spiral ganglion cells—known as cochlear synaptopathy (CS, Kujawa & Liberman, 2009)—can only be quantified via post-mortem temporal bone histology (Makary et al., 2011; Viana et al., 2015; Wu et al., 2018). Understanding how various aspects of SNHL contribute to speech recognition deficits remains a major challenge in hearing science, particularly due to the difficulties in diagnosing synaptopathy non-invasively in humans (Plack et al., 2016; Hickox et al., 2017; Kobel et al., 2017; Bramhall et al., 2019; DiNino et al., 2022). Nonetheless, studies in both animals and humans indicate that CS is a critical component of SNHL, as it often progresses with age (Sergeyenko et al., 2013; Parthasarathy & Kujawa, 2018; Wu et al., 2018) and occurs prior to OHC damage following noise exposure (Kujawa & Liberman, 2009; Fernandez et al., 2015). This suggests that the prevalence of synaptopathy may significantly exceed the World Health Organization’s estimate of 5.3% of the global population suffering from disabling hearing loss as diagnosed using the audiogram (Stevens et al., 2013; WHO, 2019).

Cochlear synaptopathy is thought to contribute to reduced speech intel-ligibility in individuals with otherwise normal audiograms (Mepani et al., 2021) due to its association with compromised supra-threshold temporal envelope (TENV) coding (Bharadwaj et al., 2014). This connection may help explain why hearing sensitivity alone is not a reliable predictor of individual speech-intelligibility scores (Festen & Plomp, 1983; Papakonstantinou et al., 2011). In animals with histologically verified CS, compromised TENV coding can be measured using envelope-following responses (EFRs, Parthasarathy et al., 2014; Shaheen et al., 2015; Parthasarathy & Kujawa, 2018). The EFR is a non-invasive auditory-evoked potential (AEP) that reflects the phase-locked neural response of a population of peripheral and brainstem neurons to a stimulus envelope (Kraus et al., 2017). Its frequency-domain response displays peaks that correspond to both the stimulus modulation frequency and its harmonics. Because EFRs can be reliably recorded in both animals and humans, their spectral magnitude is proposed as a non-invasive marker of synaptopathy that can be used across species (e.g., Shaheen et al., 2015).

To develop effective treatments for SNHL, it is essential to determine whether CS affects speech intelligibility. This goal has led to numerous studies exploring this potential relationship in humans, as summarized by DiNino et al. (2022). However, the findings have been mixed and often inconclusive. For example, while the amplitude of wave I in the auditory brainstem response (ABR), a marker of CS in animal studies (Kujawa & Liberman, 2009; Möhrle et al., 2016), has been shown to predict speech intelligibility in some research (Bramhall et al., 2015; Liberman et al., 2016), it has not produced consistent results across all studies (Johannesen et al., 2019; Prendergast et al., 2017). Similarly, studies based on EFR markers of CS also show variability; some indicate that certain EFR markers can predict speech intelligibility in individuals with normal hearing (Mepani et al., 2021), while others do not support this finding (Guest et al., 2018; Grose et al., 2017).

Several factors may contribute to the inconsistent outcomes across studies:

- **Individual differences**: Variations in the absolute strength of the ABR and EFR can reflect non-hearing-related factors in humans, such as head size (Trune et al., 1988; Mitchell et al., 1989; Plack et al., 2016). This suggests the need for a relative metric design to improve their sensitivity to SNHL aspects (Bharadwaj et al., 2015; Mehraei et al., 2016; Hickox et al., 2017; Le Prell, 2019).
- **Mixed signals**: Both ABR and EFR can indicate a combination of CS and OHC deficits (Verhulst et al., 2016b; Garrett & Verhulst, 2019; Van Der Biest et al., 2023). Without histopathological data, it is challenging to interpret these responses specifically in terms of synaptopathy.
- **Sensitivity of recording paradigms**: The recording methods developed in animal studies may not be sufficiently sensitive for use in humans, potentially overlooking cases of CS (Hickox et al., 2017; Bramhall et al., 2019).
- **Broadband nature of speech**: While speech is a broadband signal, the EFR primarily reflects auditory TENV coding mechanisms associated with ANF activity at higher cochlear frequencies, above the phase-locking limit (Joris & Yin, 1992; Verschooten et al., 2015; Henry et al., 2016). Speech recognition also relies on frequency content below this limit, which is encoded as temporal fine structure (TFS) information by the ANFs. This TFS information serves as a critical perceptual cue (e.g. Lorenzi et al., 2006; Hopkins et al., 2008; Hopkins & Moore, 2010; Henry et al., 2016; Mai & Howell, 2023; Borjigin & Bharadwaj, 2023) and is likely not reflected well in the EFR. Thus, given that conventional EFR markers mainly assess TENV coding and its impairments, the lack of correlation between EFR (driven by TENV) and speech recognition (which involves both TFS and TENV coding) does not necessarily mean that CS does not affect speech intelligibility. It may also indicate that distinct mechanisms govern each metric independently.

This study aims to clarify factors that may have complicated the interpretation of the relationship between speech perception and AEP markers of CS. First, we focus on EFRs rather than ABRs. This choice is based on previous model simulations and data, which suggest that OHC deficits have a more pronounced effect on the ABR amplitude compared to the EFR amplitude (Verhulst et al., 2016a; Garrett & Verhulst, 2019; Vasilkov et al., 2021). Second, auditory model simulations have shown that optimizing the EFR stimulus envelope to be rectangular, rather than sinusoidal, can improve the EFR marker’s sensitivity to CS (Vasilkov et al., 2021).

Building on these findings, our study first assesses the sensitivity of an optimized EFR marker for CS in a Budgerigar model with kainic acid-induced ANF damage. We then validate its effectiveness in detecting individual differences in CS compared to conventional EFR markers. We proceed to examine the correlation between this EFR marker and speech intelligibility using both low-pass (LP) and high-pass (HP) filtered speech materials in human subjects. The idea behind using filtered speech stimuli is that the listener would rely on specific auditory processing cues associated with low or high-frequency hearing. In this context, the perception of the LP condition is predominantly based on available temporal fine structure (TFS) cues (Lorenzi et al., 2006), whereas the HP condition mostly relies on temporal envelope (TENV) cues. It is well known that auditory-nerve phase-locking to TFS declines with increasing frequency (i.e., 1.4 kHz in humans based on interaural-time-difference sensitivity; Joris & Verschooten, 2013), and that for frequencies beyond this limit, the auditory system can only rely on TENV processing. Our hypothesis is that the EFR marker offers more accurate predictions of speech recognition thresholds when both measures depend on high-frequency TENV mechanisms and their associated impairments. To delineate the individual contributions of CS and hearing sensitivity to speech intelligibility, we investigate the connections between objective markers of hearing and speech intelligibility in three groups: young individuals with normal hearing, and two age-matched groups of older participants: one with normal audiograms and the other with impaired audiograms. We presume that the older participants may experience age-related CS (Wu et al., 2018), potentially compounded by additional pathologies affecting OHCs.

## Materials and Methods

### Study Participants

Three participant groups were recruited using age and audiometric puretone thresholds as the selection criteria: yNH, oNH and oHI. The younger (y) subjects were between 20 and 30 years old, and older (o, OLD) subjects had ages between 60 and 70. Normal-hearing (NH) subjects had audiometric thresholds less than 20 dB HL for frequencies up to 4 kHz. Hearing-impaired (HI) subjects had sloping audiograms and thresholds that surpassed 20 dB HL at least once below 4 kHz. The grouping criterion did not account for potential individual variations in the degree of synaptopathy; this was an unknown variable at the start of the study. Using these criteria, fifteen young normal-hearing (yNH: 24.5 *±* 2.2 y/o, 8 females), 15 older normal-hearing (oNH: 64.2 *±* 1.9 y/o, 7 females) and 14 older hearing-impaired (oHI: 65.2 *±* 1.7 y/o, 7 females) subjects participated. There were no significant age differences between the oNH and oHI participant groups (p*>*.05). Otoscopy was performed prior to data collection to ensure that participants had no obstructions or other visible outer or middle ear pathologies. Aside from the audiogram and otoscopy, all tests were performed monaurally on the ear with the best audiometric thresholds. The experiments were approved by the ethics committee of the University of Oldenburg. Participants gave written informed consent and were paid for their participation.

### Behavioral and Physiological Markers of Hearing Sensitivity

We adopted two measures to quantify hearing sensitivity: a standard clinical pure-tone audiogram to assess the behavioral hearing sensitivity, and distortion-product otoacoustic emission (DPOAE) thresholds (TH_DP_) to assess the OHC integrity more directly through changes in OHC-driven ear canal pressure. We collected DPOAEs at 4 kHz to quantify OHC-damage in the same frequency region as targeted by the EFR marker of CS.

DPOAE stimuli were presented over ER-2 speakers (Etymotic Research) using foam ear tips and DPOAEs were recorded using the ER10B+ OAE microphone system (Etymotic Research) and custom-made MATLAB scripts (Mauermann, 2013). Two pure tones (f_1_, f_2_) were simultaneously presented at a fixed f_2_/f_1_ ratio of 1.2 using a primary frequency-sweep method (Long et al., 2008). Frequencies were exponentially swept up (2 s/octave) over a 1/3*^rd^* octave range around the geometric mean of 4 kHz. Primary level L_1_ followed the Scissors paradigm (L_1_=0.4 L_2_+39; Kummer et al., 1998) given a primary L_2_ of 30-60 dB SPL in steps of 6 dB (in oHI listeners, L_2_ of 66 and 72 dB SPL were additionally collected). The distortion component (L_DC_) was extracted using a sharp 2-Hz-wide least-squares-fit filter and the center frequency of the measured frequency range was used to construct L_DC_ growth functions. Individual L_DC_ data points and their standard deviations were used in a bootstrapping procedure to fit an adapted cubic function through the L_DC_ datapoints as described in Verhulst et al. (2016a). L_DC_ growth functions typically increase monotonically or saturate with increasing L_2_ (Mauermann & Kollmeier, 2004; Abdala et al., 2021). We thus constrained our bootstrapping procedure to only include random L_DC_ draws (i.e. from within the confidence interval of each mean L_DC_ measurement point) to impose monotonous growth in each L_DC_ (L_2_,b) bootstrap run. We used an automated algorithm which eliminated adjacent data points at either end of the growth function (never intermediate points) that compromised monotonicity. TH_DP_ was determined in each bootstrap run as the L_2_ at which the extrapolated fitting curve reached a level of -25 dB SPL (Neely et al., 2009). This bootstrapping procedure yielded the median threshold (TH_DP_) and its standard deviation. TH_DP_ of two yNH participants with reasonable growth functions but very low thresholds (-30.9, -5.4 dB SPL) were set to 1.5-times the interquartile range of the yNH TH_DP_ threshold distribution (-4.3 dB) to avoid high-leverage data-points. TH_A_ and TH_DP_ correlated strongly at 4 kHz (*ρ*(44) = 0.84; p *<* 0.05), corroborating earlier observations (Boege & Janssen, 2002) and confirming that both metrics assess hearing sensitivity.

Audiometric thresholds (TH_A_s) were collected for frequencies between 0.125-8 kHz using Sennheiser HDA200 headphones and a clinical audiometer AT900 (Auritec). Figure 1 shows audiograms of the tested ears, which were chosen based on the better of left and right audiograms. At 4 kHz, yNH_control_ participants had a mean (M*±*SD) threshold of 3.3 *±* 3.5 dB HL. The older groups had thresholds of 11.3 *±* 3.5 dB HL (oNH) and 36 *±* 8.4 dB HL (oHI) respectively.

**Figure 1:**
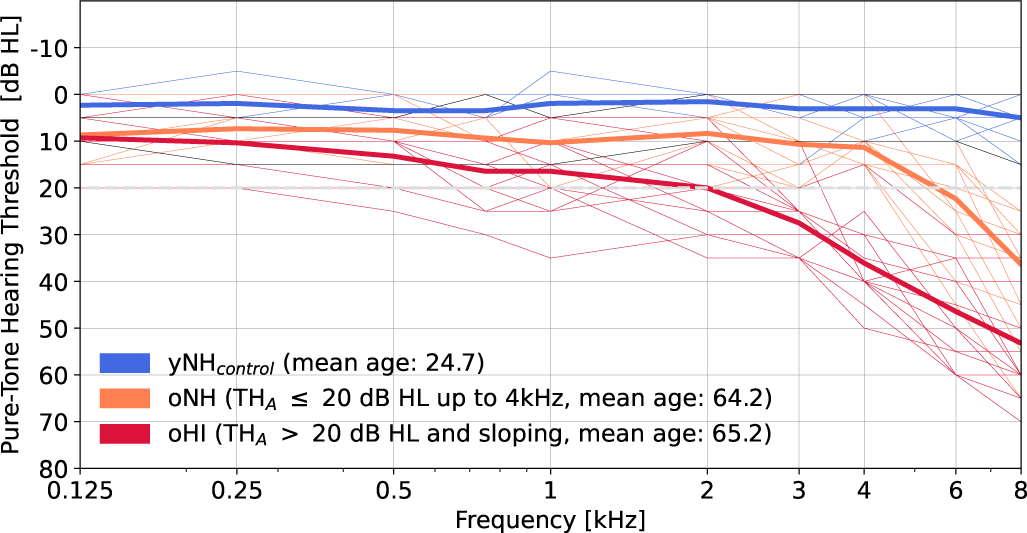
Pure-tone hearing thresholds (in dB HL) at frequencies between 0.125-8 kHz. Groups were based on audiogram and DPOAE thresholds and age. yNH_control_: young normal hearing control group (blue), oNH: old normal hearing (orange), oHI: old hearing impaired (red). Thick traces represent the group mean and thin traces represent individual audiogram profiles. Note that two subjects from the recruited NH group did not meet the TH_DP_ threshold criterion for normal hearing to be included in the control group. The audiograms of these NH subjects were indicated with thin black traces. The 20 dB HL hearing threshold (TH_A_), which was used to separate listeners into oNH or oHI subgroups, was indicated by a gray dashed curve.

### Human envelope-following-response recordings

EFRs were recorded in humans to study their relationship to speech intelligibility and in Budgerigars to assess their sensitivity to kainic-acid induced CS. Human EFRs were recorded to two amplitude-modulated pure-tone stimuli with the same carrier frequency (f = 4 kHz) and modulation frequency (f_m_ = 120 Hz, starting phase 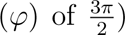. The only difference between the stimuli was their envelope shape: the SAM stimulus had a sinusoidal modulator (Eq.1), as commonly used in animal studies of CS (e.g., Shaheen et al., 2015; Parthasarathy & Kujawa, 2018) while the modulator of the second stimulus was a rectangular-wave with a duty-cycle (*τ*) of 25% (RAM; Vasilkov et al., 2021). A modulation depth (md) of 0.95 (95%, -0.45 dB re. 100%) was applied. MATLAB’s ‘square’ function was adopted to create the RAM stimulus, and both stimuli are formulated as follows:

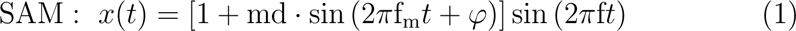

RAM : *y(t)* = [2 + 2md · *m(t)*] sin (2φ*ft*), with

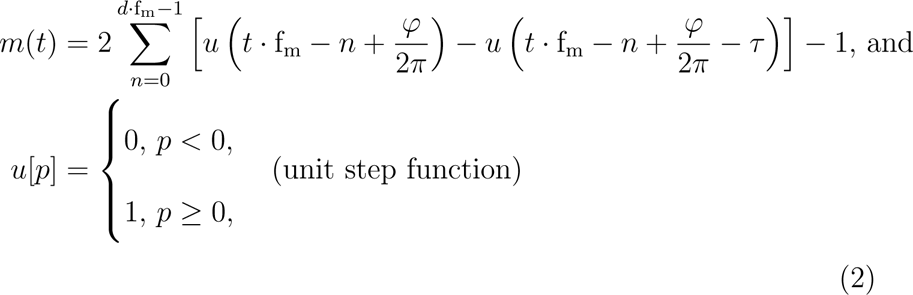

Stimuli were windowed using a 2.5% tapered-cosine window, had a duration (d) of 0.4 seconds and were repeated 1000 times each (500 per polarity). The inter-stimulus interval consisted of a uniformly distributed random silence jitter (100 ms *±* 10 ms). The SAM tone was presented at 70 dB SPL and the RAM stimulus at the same peak-to-peak amplitude, which corresponded to 68 dB SPL. A previous study compared a broad range of possible ABR and EFR markers of CS (Vasilkov et al., 2021), and we selected the most promising RAM stimulus for use in the present study. Stimuli were generated in MATLAB (R2015b) at a sampling rate of 48 kHz and calibrated using a Brüel & Kjær ear simulator type 4157 for insert earphones. A Fireface UCX sound card (RME) and TDT-HB7 headphone driver (Tucker-Davis) were used to drive the ER-2 insert earphones (Etymotic Research) using the open-source portaudio playrec ASIO codec (Humphrey, 2008). Stimuli were presented monaurally to the test ear.

Recordings took place in a double-walled electrically-shielded measurement booth (IAC acoustics) and participants sat in a reclining chair while watching a silent movie. EEG signals were recorded using a 64-channel cap with equidistant electrode spacing (Easycap) and active Biosemi Ag/AgCl electrodes were connected to a Biosemi amplifier. A sampling rate of 16384 Hz and 24-bit analog-to-digital conversion were used to store the raw data traces. A common-mode-sense (CMS) active electrode was placed on the fronto-central midline and a driven-right-leg (DRL) passive electrode was placed on the tip of the nose. Reference electrodes were placed on each earlobe. Electrode offsets (DC values of the CMS signal) were kept below 25 mV.

Raw EEG recordings were extracted in Python (version 2.7.10 | Anaconda 2.3.0 (64-bit), www.python.org) and MNE-Python (version 0.9.0; Gramfort et al., 2013, 2014) and all EEG recording channels were re-referenced to the offline-averaged earlobe electrodes. Data were epoched in 400 ms windows starting from the stimulus onset and baseline corrected by the average amplitude per epoch. We only present results from the vertex channel (Cz) in this study which is known to yield good signal strength for subcortical auditory sources (Picton, 2010). Signal processing was performed in MATLAB (R2014b). The EFR estimates for each stimulus condition and participant were computed based on the energy at the modulation frequency and its first four harmonics (h_0_-h_4_ = *k ×* f*_m_*, *k*=[1..5]) to account for all envelope-related energy in the EEG signal (Vasilkov et al., 2021). Equation 3 was used in a bootstrap routine to obtain the mean EFR amplitude and corresponding standard deviation (Zhu et al., 2013):

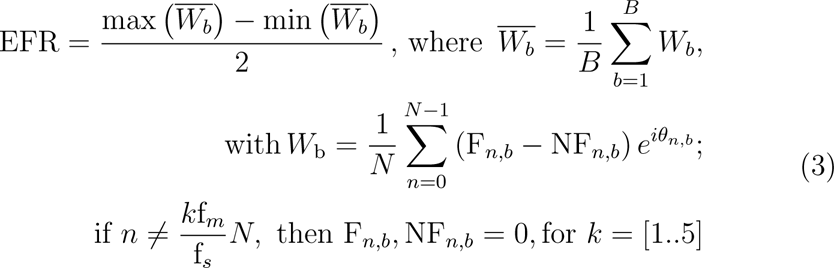

where *N* corresponds to the number of frequency bins in the magnitude spectrum. First, a mean spectral estimate of the EEG recording for each frequency component (n) was computed by averaging the complex discrete Fourier transform values of 1000 randomly drawn epochs using 500 epochs of each polarity (with replacement) in each bootstrap run (b; F_n,b_). Epochs were windowed using a 2% tapered-cosine window before the frequency-domain transformation. The electrophysiological noise floor (NF_n,b_) at frequencies h_0_-h_4_ was computed as the average magnitude of the ten frequency-bins surrounding the respective frequency (five bins each side). The noise-floor estimates were then subtracted from the signal components h_0_-h_4_, to yield peak-to-noise floor (PtN_n,b_) magnitude estimates. Figure 2A (left panels) illustrate the mean magnitude spectra and noise floor estimates for an example human recording. All frequency components apart from the harmonic frequencies (h_0_-h_4_) were removed from the noise-floor-corrected spectrum before it was transformed back to the time domain using the inverse discrete Fourier transform and the original phase information (*θ*_n,b_) of the harmonic frequencies. This procedure was repeated b=200 times to yield 200 reconstructed time-domain estimates of the EFR waveform (W_b_). The W_b_ waveforms were then averaged and the EFR was defined as half the peak-to-peak amplitude of the averaged reconstructed time-domain waveform. Figure 2A (right panels) shows the reconstructed EFR time-domain waveform for an example human recording and compares it against the original, filtered EEG signal on which the reconstruction procedure was applied. The metric defined in Eq.3 corresponds to the EFR peak-to-noise-floor amplitude and is further referred to as the EFR amplitude (in *µ*V), or the EFR marker.

**Figure 2:**
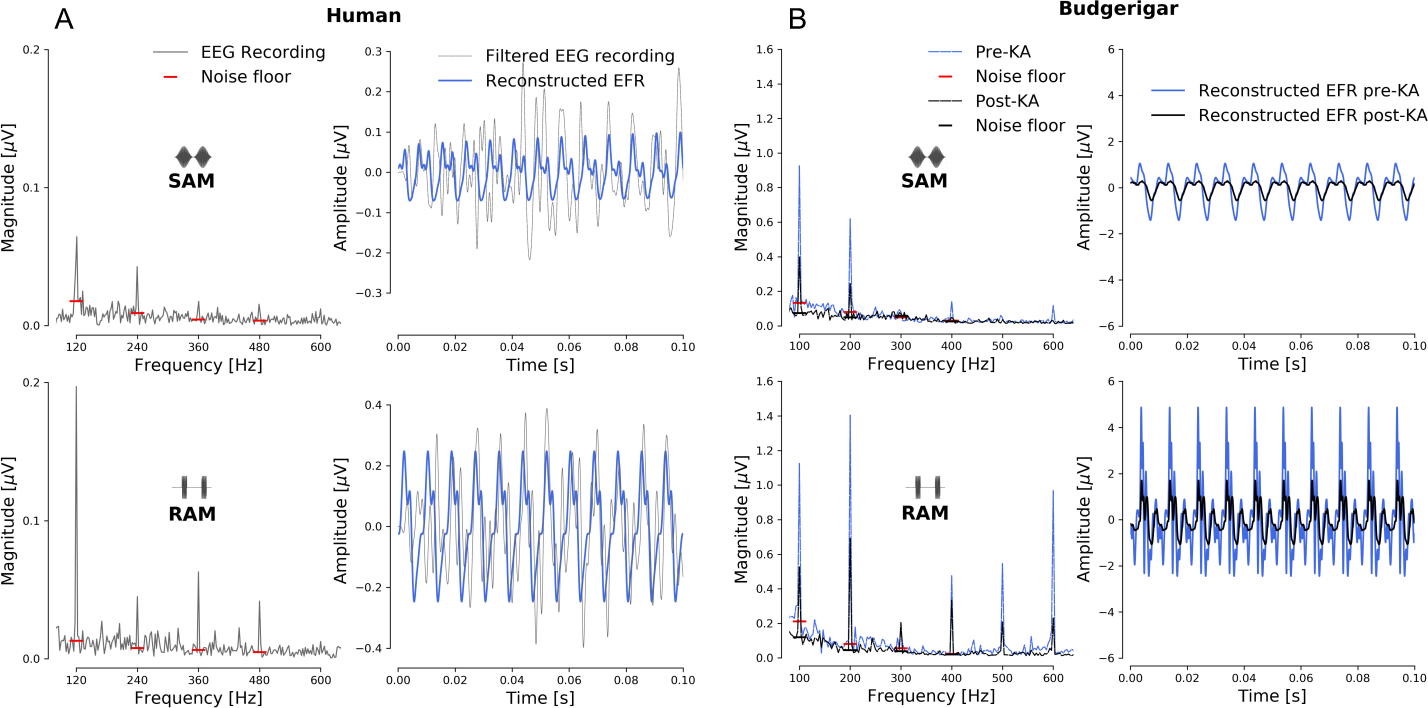
Comparison between human (panel A) and Budgerigar (panel B) EFR recordings and analysis procedures for single subjects. For each species, the top row corresponds to the SAM stimulus and the bottom row to the RAM stimulus. The modulation frequencies and depths were 120 Hz (95% md) and 100 Hz (100% md) for humans and budgerigars, respectively. Carrier frequencies were 4 kHz (68 dB SPL RAM, 70 dB SPL SAM) and 2.83 kHz (75 dB SPL), respectively. Time-domain responses (right panels) show the filtered EEG recording along with the reconstructed time-domain waveform that was based on five frequency components (h_0_-h_4_) and their respective phases. The EFR amplitude (or, EFR marker) was extracted from the reconstructed time-domain EFR following Eq.3, which was based on an iFFT of the noise-floor-corrected mean EFR magnitude (left panels). The budgerigar recordings (panel B) are shown for the same animal before or after kainic-acid (KA) administration. Post-KA spectral peaks and reconstructed EFRs were smaller than pre-KA EFRs.

### Budgerigar EFR recordings and kainic-acid induced synaptopathy

EFRs were also recorded in eighteen young adult Budgerigars (Melopsittacus undulatus), approximately two years of age, before and/or after induction of CS using kainic acid (KA). The Budgerigar is a small parrot species with human-like behavioral sensitivity to many simple and complex sounds in operant conditioning studies (Dent et al., 2000; Dooling et al., 2000), and was selected based on its use in ongoing behavioral studies of CS (Wong et al., 2019). All procedures were performed at the University of Rochester and approved by the University Committee on Animal Resources. Three animals were tested before and after KA administration and monitored over time, whereas the others in the cohort either belonged to a control-only (n=11) or KA-only group (n=4). KA is a glutamate analog that damages cochlear afferent synapses between hair cells and auditory-nerve fibers through excitotoxicity in mammals and birds (Bledsoe et al., 1981; Pujol et al., 1985; Zheng et al., 1997; Sun et al., 2001). In Budgerigars, bilateral cochlear infusion of KA has been shown to permanently reduce ABR wave-I amplitude by up to 70% without impacting behavioral audiometric thresholds or DPOAEs generated by sensory hair cells (Henry & Abrams, 2018; Wong et al., 2019). We used the methods described in Henry & Abrams (2018) and Wong et al. (2019) to induce synaptopathy in Budgerigars. Briefly, animals were anesthetized with ketamine (5-6 mg/kg) and dexmedetomidine (0.1 mg/kg; subcutaneous bolus injection) and placed in a custom stereotaxic device. Ketamine/dexmedetomidine were administered subcutaneously for anesthetic induction, but anesthesia was maintained throughout the surgery (approximately 1-2 hours) by a continuous anesthetic infusion pump (Razel Scientific; Fairfax, VT, USA). The middle-ear space was accessed surgically using a posterior approach to expose the basal prominence of the cochlear duct, where a 0.15-mm diameter cochleostomy was made using gentle rotating pressure on a small manual drill. Thereafter, 2.5 *µ*L of 2-mM KA (Abcam ab 144490; Cambridge, UK) in hanks balanced salt solution (Sigma-Aldrich H8264; St. Louis, MO, USA) was infused into cochleostomy over 90 seconds using a microinjection needle. Compound action potentials (CAPs) of the auditory nerve were recorded before and after infusion in response to clicks. Excitotoxic synaptic injury was confirmed by observing *>*90% CAP reduction within 10-20 minutes following KA exposure. The left and right ears were treated with KA during different surgical procedures four weeks apart to minimize operating time and to avoid excessive anesthetic exposure. DPOAEs were recorded using a swept-tone paradigm (see Wong et al., 2019) before and after surgeries to confirm no adverse impact of the procedures on sensory hair cells. Prior to KA exposure, wave-I amplitude of the ABR in response to 90-dB peSPL clicks was 24.56 *±* 2.31 *µ*V in animal K20 and 23.69 *±* 2.60 *µ*V in animal K25 (M *±* SD). Reduction of wave I, based on ABRs recorded four or more weeks post KA (during the steady-state period), was 68.5% in animal K20 and 64.9% in animal K25. EFRs were measured at multiple points before KA exposure and at four time points after the second infusion. Repeated measurements assessed within-subject variability of responses, since synaptic injury remains relatively stable after one month following KA exposure (Sun et al., 2000; Henry & Abrams, 2018; Wong et al., 2019). Anesthesia was performed as described above using ketamine and dexmedetomidine (for recording sessions, anesthesia was only administered subcutaneously), and body temperature was maintained in the normal range for this species of 39-41 degrees C.

Stimuli were generated in MATLAB (The MathWorks, Natick, MA, USA) at a sampling frequency of 50 kHz. Stimuli were SAM and RAM tones presented at 75 dB SPL with 10-ms cosine-squared onset and offset ramps, 300-ms duration, and a 130-ms silent interval between successive stimuli. Carrier frequency and modulation frequency were 2830 Hz and 100 Hz, respectively, and the polarity of the carrier signal was alternated between stimulus repetitions. Depth of modulation was 100%, and the duty-cycle for RAM modulation was fixed at 25%. Stimuli were converted to analog using a data acquisition card (PCIe-6251; National Instruments, Austin, TX, USA), which also digitized response activity using the same internal clock. Stimuli were presented free-field through a loudspeaker (MC60; Polk Audio, Baltimore, MD, USA) positioned 20 cm from the animals head in the dorsal direction (the rostral surface of the head faced downward in the stereotaxic apparatus; thus the loudspeaker and animal were located in the same horizontal plane). Level was controlled digitally (by scaling stimulus waveforms in MATLAB) and by up to 60 dB of analog attenuation applied by a programmable attenuator (PA5; Tucker Davis Technologies, Alachua, FL, USA). Calibration was based on the output of a <u>^1^</u> “ precision microphone (model 4938; Brüel and Kjær, Marlborough, MA USA) in response to pure tones.

Electrophysiological activity was recorded differentially between a stain-less steel electrode implanted at the vertex (M0.6 threaded machine screw; advanced through the skull to the level of the dura) and a platinum needle electrode (model F-E2; Natus Manufacturing, Gort, Co. Galway, Ireland) inserted at the base of the skull near the nape of the neck. A second needle electrode in the animal’s back served as ground. Activity was amplified by a factor of 50,000 and filtered from 30-10,000 Hz (P511; Grass Instruments, West Warwick, RI USA) prior to sampling (50 kHz) by the data acquisition card. Responses to 300 repetitions of the same stimulus (including both polarities) were averaged to produce each EFR waveform and amplitude, following the procedure described for human EFR recordings in Eq.3. The Budgerigar panels in Fig.2B (left panels) depict the mean EFR spectral magnitudes and noise floor estimates for the SAM and RAM stimulus for a Budgerigar that was monitored longitudinally. Spectra and reconstructed EFR waveforms are shown both before and after KA administration and demonstrate smaller EFR waveforms after ototoxic-induced CS in this species (Fig.2B, right panels).

### Human Speech Reception Thresholds (SRTs)

Speech intelligibility was assessed by applying a standard German Matrix sentence test which determines the speech reception threshold (SRT) in quiet or in a fixed-level noise background (Matrix-test or OLSA; Wagener et al., 1999; Brand & Kollmeier, 2002). The OLSA test consists of 5-word sentences (male speaker: name - verb - numeral - adjective - object) using a corpus of 50 possible words. The speech-shaped background noise was generated from the sentences and matched the long-term spectrum of the speech material. The SRT for 50% correctly identified words (i.e., SRT_50_) was determined using a 1-up/1-down adaptive procedure with varying step size based on word scoring (20 sentences per condition).

The SRT_50_ was determined in quiet (SiQ) and noise (SiN) for unfiltered/broadband (BB), low-pass filtered (LP) and high-pass filtered (HP) audio. The speech and noise signals in the LP and HP conditions were generated by applying a 1024^th^ order FIR filter with respective cut-off frequencies of 1.5 and 1.65 kHz to the OLSA test material (i.e., the BB condition). Since the adopted EFR marker predominantly captures high-frequency TENV processing, it is logical to incorporate a speech condition that similarly depends on cochlear TENV processing, such as the SRT_HP_.

Both SiQ and SiN conditions were included in our study because we anticipate that speech processing in the presence of a fixed-level background noise may be more adversely affected by CS. Although both hearing sensitivity (Plomp, 1986) and CS-compromised TENV processing (Shaheen et al., 2015; Parthasarathy & Kujawa, 2018) can impact SiQ processing, we believe that the SiN condition may be further compromised by reduced coding redundancy due to AN deafferentation (Lopez-Poveda & Barrios, 2013). In the SiQ conditions, the speech level was varied and the dB SPL at which the 50% correct threshold was reached, was reported. The initial speech level in the SiN test was 70 dB SPL and the noise level was kept fixed at 70 dB SPL while the speech level varied adaptively to yield the SRT. The six conditions (3 SiQ, 3 SiN) were presented in pseudo-random order. Participants completed three training runs in which a SiN_BB_ with a fixed SNR of 5 dB was followed by a regular SiN_BB_ condition with SRT tracking for training purposes. All possible words were displayed on the screen during those runs to familiarize participants with the stimulus material. The third training run was a SiN_HP_ condition with SRT tracking but without visual aid. During the experiment, answers were logged by the experimenter and no visual feedback was provided. Measurements were conducted in a double-walled sound-insulated booth using Sennheiser HDA200 headphones in combination with the Earbox Highpower ear 3.0 sound card (Auritec) and the Oldenburg Measurement Platform (HörTech gGmbH). The setup was calibrated using an artificial ear type 4153, microphone type 4134, preamplifier type 2669 and sound level meter type 2610 (Brüel & Kjær).

### Statistical and post-hoc analysis

To disentangle the effects of CS and hearing sensitivity on the EFR markers and SRTs, we performed group statistics, as well as a multiple regression analysis using the pooled data. TH_A_s and TH_DP_s were used as criteria to assign listeners to specific groups. Aside from their normal audiometric thresholds, all subjects in the yNH_control_ group were ensured to have a TH_DP, 4 kHz_ *≤* 25 dB SPL, to minimize the risk of OHC damage in this group. As a result, two participants from the original yNH group were excluded from the yNH_control_ group based on their TH_DP, 4 kHz_. Their datapoints were, however, included in the multiple regression analysis across all individuals, and were identified with distinct markers in the corresponding regression figures. We performed a group analysis using the ‘aov group’ function from the R programming environment (R Core Team, 2019), and investigated main effects of age and hearing sensitivity using unpaired Students’ t-tests between the yNH_control_ (n=13) and oNH (n=15) groups, and between the oNH and oHI (n=14) groups, respectively. The ‘SciPy’ python package for scientific computing (Oliphant, 2007; Millman & Aivazis, 2011) and ‘stats.ttest ind’ function was used for this purpose. p-values for multiple comparisons were Bonferroni adjusted to control for the family-wise error rate. The applied correction factors are given in the Results section.

We examined linear correlations between the EFR and SRT on the pooled data (n=44) using correlation coefficients calculated using the ‘SciPy’ python package and ‘stats’ package. All correlations refer to the Pearson correlation coefficient (r) if both variables were normally-distributed (Shapiro-Wilk test), otherwise the Spearman’s rank correlation coefficient (*ρ*) was reported. These correlations were further investigated using using the ‘sklearn’ linear regression functions in Python to analyse the residuals, and using multiple regression models to verify the contribution of age and hearing sensitivity (’lsmeans’ package in R Russell, 2016). Additionally, we performed commonality analysis using the ‘yhat’ package (Nimon et al., 2008). Commonality analysis combines linear regressions on the dependent variable and allows for the decomposition of the explained variance (R^2^) of the linear predictors into subcomponents explained by the unique and the common/shared variance of predictors and all their possible combinations (Newton & Spurrell, 1967). This technique also works in the presence of multicollinearity (Ray-Mukherjee et al., 2014).

Because hearing sensitivity, as assessed behaviorally using pure-tone thresholds is similar, but not necessarily identical, to OHC sensitivity assessed through DPOAE thresholds (Boege & Janssen, 2002), we performed additional post-hoc statistics using the TH_DP, 4 kHz_ as the measure of hearing sensitivity. For the group-based analyses, we divided the cohort into groups with either normal or impaired DPOAE thresholds, reflecting TH_DP, 4 kHz_ *≤* or *>* 25 dB SPL, respectively. We also considered the pooled older group (OLD, n=29) when investigating main effects of age-related CS in comparison to the yNH_control_ group, and the pooled normal-hearing group (NH, n=28) when investigating main effects of hearing sensitivity in comparison to the oHI group.

## Results

### Envelope-following-response sensitivity to cochlear synaptopathy

Panel B in Figure 2 shows the effect of KA on Budgerigar SAM and RAM EFR magnitude spectra (left) and reconstructed waveforms (right). Energy at the modulation frequency of the stimulus and its harmonics was reduced after KA administration, leading to an overall reduction in the reconstructed EFR waveform and its corresponding amplitude. EFR amplitude reductions occurred consistently across the three longitudinally-monitored Budgerigars (Fig.3A), and was attributed to a histology-verified reduction in auditorynerve peripheral axons and cell bodies (Fig.3B). The histology confirms that KA introduces cochlear synaptopathy in Budgerigars, and the additionally performed DPOAE analysis in Wang et al. (2023) and Wilson et al. (2021) furthermore demonstrates the selectivity of KA to AN synapses and cells without damaging the OHCs. As Fig.3A depicts, EFR amplitude reductions occur instantly after KA administration, and recover slightly to an overall reduced amplitude over the following weeks.

**Figure 3:**
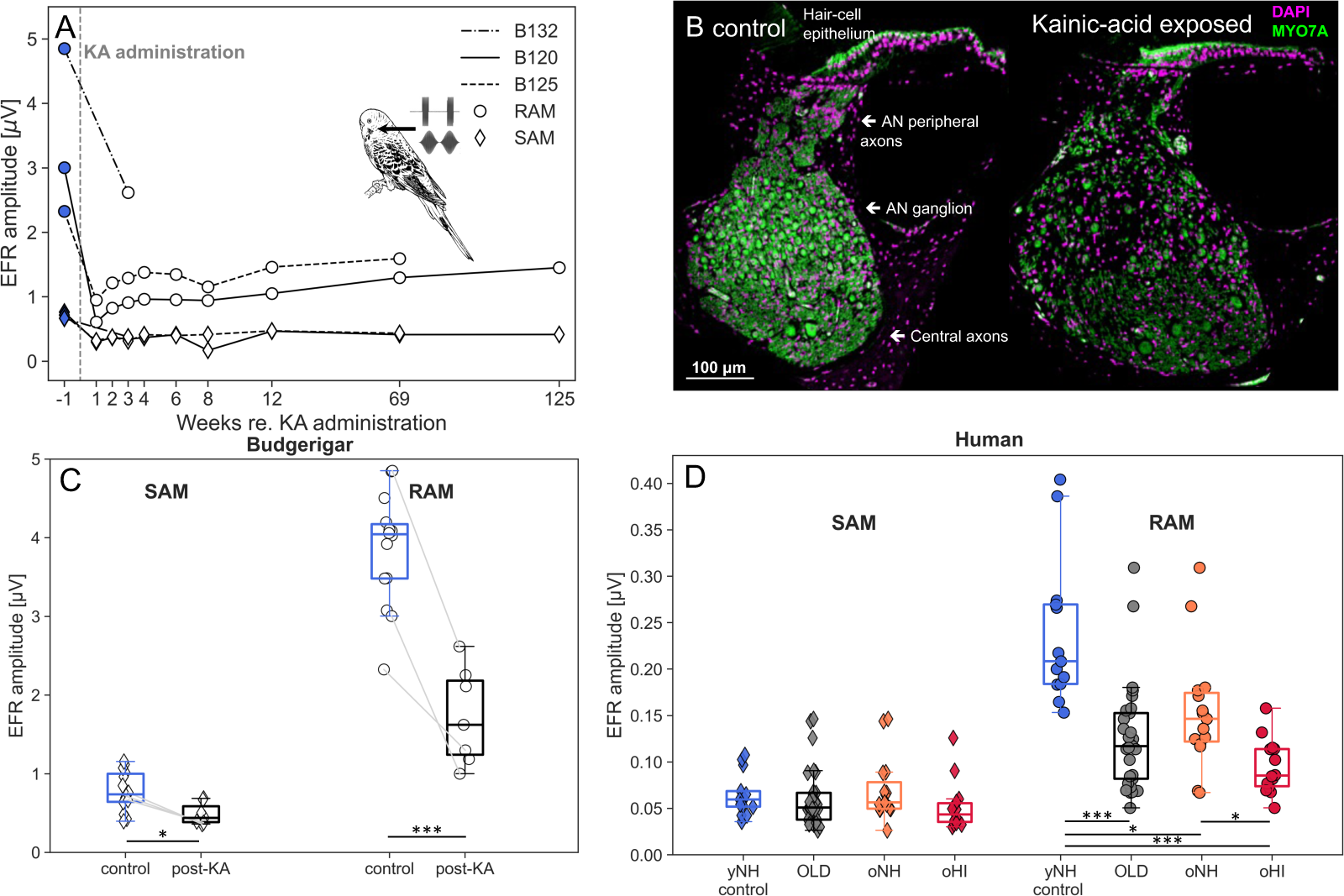
**A** EFR amplitudes from three Budgerigars (B132, B120, B125) before or several weeks after administration of kainic acid (KA). **B** Representative cross sections of the Budgerigar cochlea from a control ear (left) and from an ear exposed to 1-mM KA solution (12 weeks post exposure; right). Sections are stained for DAPI and Myosin 7A, and are from the location 50-60% of the distance from the apex to the base (≈2-kHz cochlear frequency; see Wang et al., 2023). KA exposure causes marked reduction of auditorynerve (AN) peripheral axons and cell bodies in the AN ganglion, without impacting the hair-cell epithelium. **C** Boxplots and individual data points of the Budgerigar SAM and RAM EFR amplitudes before or after KA-administration. Connected lines correspond to data from the same animal. To account for both paired and independent data in the Budgerigar sample, we performed a partially overlapping samples t-test and reported significance as: (*) p*<*.05, (**) p *<* .01 and (***) p *<* .001. **D** Boxplots and individual data points of human SAM and RAM EFR amplitudes. Data is show for the different test groups based on age and TH_A_s: yNH_control_, oNH and oHI, as well as for the pooled OLD group (oNH+oHI). Independent samples t-tests were performed between all conditions for the SAM and RAM conditions separately, and significance was reported as: (*) p*<*.05, (**) p *<* .01 and (***) p *<* .001 after applying a Bonferroni correction of 6.

The post-KA EFR amplitudes shown in Fig.3C correspond to average EFR amplitudes over the different post-KA measurement time points for each animal and are compared to control (pre-KA or non-KA) EFR amplitudes. Connected lines refer to data points stemming from the same Budgerigar, and SAM and RAM EFRs were recorded during the same session. Other points show data from control animals (n=11) or post-KA exposure animals (n=4; i.e., animals for which pre-exposure EFRs were not recorded). To account for the mix of paired and independent observations in the sample, we conducted a partially overlapping samples t-test (Derrick, 2017), which revealed a significant decrease in EFR amplitude from the pre-KA condition (M = 0.78, SD = 0.245) to the post-KA condition (M = 0.49, SD = 0.127) for the SAM stimulus (t(19.89) = 2.75, p =.012). The RAM condition showed an even larger significant reduction from pre-KA (M = 3.86, SD = 0.689) to post-KA (M = 1.73, SD = 0.565, t(19.89) = 7.611, p *<*.001). The latter observation relates to the almost five-times larger pre-KA RAM amplitudes. These recordings confirm the positive effect the stimulus envelope has on the EFR signal-to-noise ratio. Single-unit AN recordings show more synchronized AN responses to faster-rising stimulus envelopes (Dreyer & Delgutte, 2006), and the AN and EFR model simulations performed in Vasilkov et al. (2021) show that this effect also impacts the neural generators of the EFRs. We conclude that the RAM EFR is a selective non-invasive marker of cochlear synaptopathy in Budgerigar and that the RAM EFR has a better sensitivity over the SAM EFR in identifying individual differences in CS. This renders the RAM EFR a suitable candidate for use in human studies for whom the intrinsic EEG signal-to-noise ratio is inherently smaller than in research animals.

### Human EFR recordings: age-related deficits

Figure 3D depicts human SAM and RAM EFRs for yNH_control_ and older subjects with or without impaired audiograms. The human EFR amplitudes are in agreement with both model predictions (Vasilkov et al., 2021) and Budgerigar findings (panel C) in showing overall 3.7 times larger RAM (M=0.239, SD=0.076) than SAM (M=0.065, SD = 0.022) amplitudes (V) in the yNH_control_ group. Compared to the yNH_control_ group, older subjects showed reductions in the amplitude of the SAM and RAM EFRs by 7% and 47%, respectively. The mean amplitudes for the older group were M = 0.061 (SD = 0.031) for SAM and M = 0.126 (SD = 0.057) for RAM. There were no significant differences between the yNH_control_ group and older listeners for the SAM EFR (p*>*.05). However, RAM EFR amplitudes were significantly reduced in the older group compared to the yNH control group (t(40)=5.17, p*<*.001). Figure 3D visualises this trend, and furthermore shows that the RAM EFRs of the oNH and oHI subgroups are significantly smaller than those of the yNH_control_ group. Together, this supports the view that the RAM EFR is more sensitive to detecting age-related changes than the SAM EFR in humans.

Humans differ from the KA Budgerigar model of CS in that it cannot be ruled out that the reductions in human EFRs reflect more than just agerelated CS. For example, our human cohort may have had other forms of SNHL (e.g., OHC damage) that could also have affected the RAM EFR marker. To investigate the potential influence of hearing sensitivity on the observed EFR reductions, the main effect of age can be considered against the main effect of hearing sensitivity. The yNH_control_ group had significantly larger RAM EFR amplitudes (M = 0.239, SD = 0.076) than the oNH group (M = 0.155, SD = 0.062, t(26) = 3.09, p = .005), and the oNH group had better EFRs than the oHI group (M = 0.095, SD = 0.028, t(27) = 3.19, p = .004). However, the main effect of age (t(40)=3.92, p*<*.001) was greater than that of TH_A_ differences in hearing sensitivity (t(27) = 3.19, p = .004). We performed an additional post-hoc regrouping using the TH_DP_ criterion of 25 dB SPL to separate cohort into those with normal or impaired OHC integrity at 4 kHz. Within the group of TH_DP_s *<* 25 dB SPL, younger subjects had significantly larger EFRs than the older listeners (t(17) = 3.9, p*<*.001), and among the older listeners, there were no significant EFR differences between those with normal or impaired TH_DP_s (t(27) = 1.1, p*>*.05). The mean TH_DP_ difference of 20.9 dB between the older subjects with normal or impaired hearing sensitivity was thus not reflected in their EFR amplitudes. Taken together, our group analyses support a predominant age-related CS interpretation of the RAM-EFR. Going further, we investigate the degree to which the RAM-EFR marker can predict speech intelligibility declines in older listeners.

### Speech reception thresholds

Individual and group speech reception thresholds (SRTs) are depicted in Fig.4 for quiet (SiQ; panel A) and stationary noise backgrounds (SiN, panel B). For each filtered condition, SRTs of the yNH_control_ group are compared to the pooled older group, as well as oNH and oHI subgroups.

**Figure 4:**
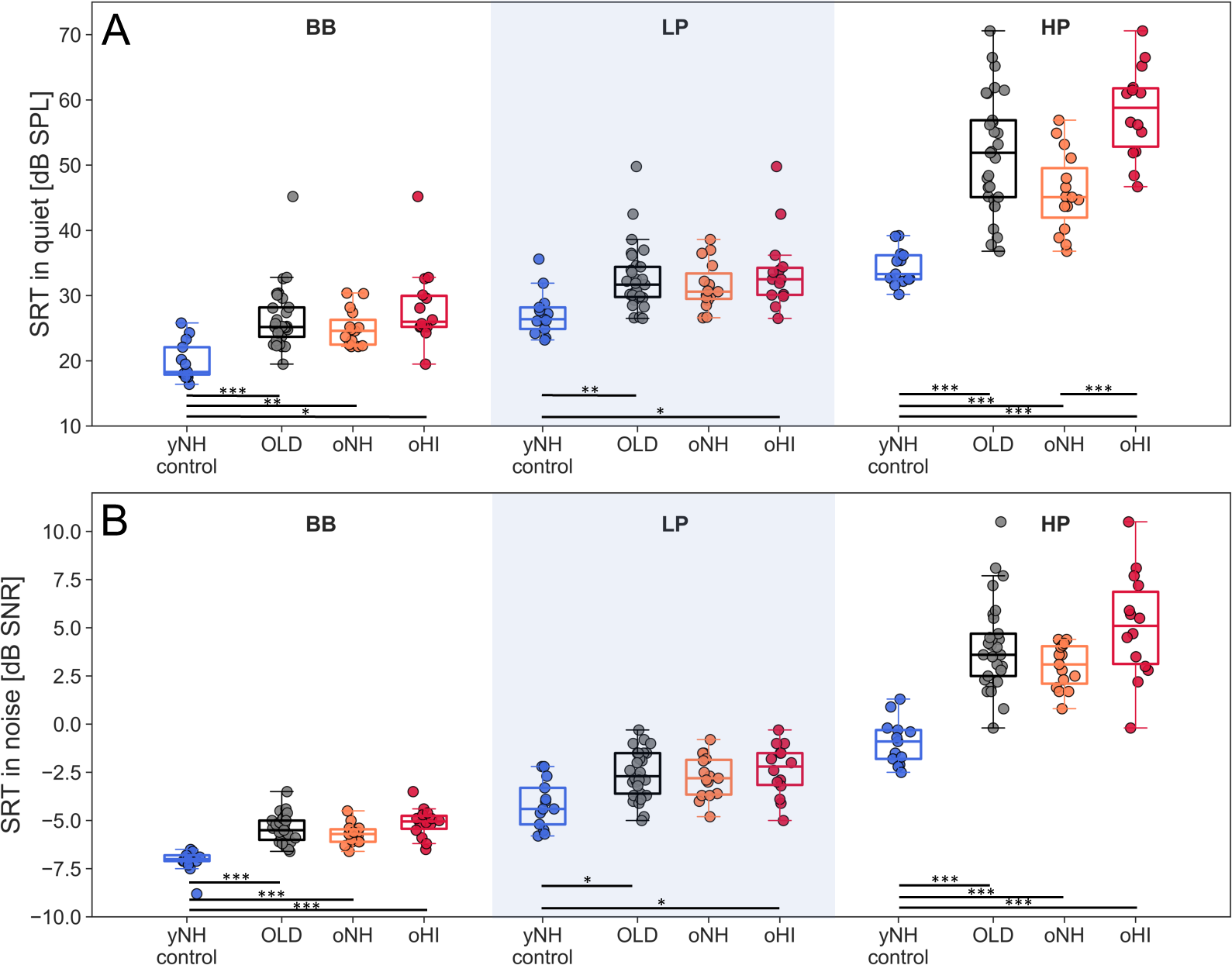
Speech reception thresholds (SRT) for the OLSA matrix sentence test presented in quiet. **(A)** and in speech-shaped noise **(B)** for three conditions: original (BB), lowpass filtered speech and noise material (f_c_=1.5 kHz; LP), high-pass filtered speech and noise material (f_c_=1.65 kHz; HP). SRTs are grouped by the selection groups (yNH_control_, oNH, oHI), as well as pooled across oNH and oHI subjects into an older group (OLD). Independent t-tests were computed between the groups in each condition for the quiet and noise conditions separately, and significant differences were indicated on the figure. The p-values were Bonferroni corrected before their significance was reported as (*) p*<*.05, (**) p *<* .01 and (***) p *<* .001. A Bonferroni correction of 18 was applied for the yNH_control_, oNH and oHI group comparisons, and of 6 for the yNH_control_ and OLD comparisons.

A two-way (3×3) mixed-design ANOVA analysis investigated the role of participant group (yNH_control_, oNH, oHI) and filter-condition (LP, HP, BB) on the SRT. Apart from significant main effects of group (SiQ: F(2,39) = 37.2, p*<*.001; SiN: F(2,39) = 34.8, p*<*.001) and filter-condition (SiQ: F(2,78) = 420.3, p*<*.001; SiN: F(2,78) = 707, p*<*.001), the interaction terms were also significant (SiQ: F(4,82) = 24.6, p*<*0.001; SiN: F(4,82) = 17.9, p*<*0.001), indicating that group SRTs were differently affected by the filtering.

Adding background noise affected the groups in the HP condition differently across the SiQ and SiN condition, while the trends observed in the BB and LP conditions remained consistent. The addition of background noise particularly impaired the oNH group in the HP condition. While the OLD subgroups performed worse than the yNH_control_ group in both SiQ and SiN conditions (t(40)=-6.8 (SiQ) and -7.3 (SiN), p*<*.001), the oNH group performed equally poorly as the oHI group in the SiN condition (p*>*.05), but not in the SiQ condition (t(27)=-5.08, p*<*0.001). This supports the view that processing TENV information in a stationary noise background is more affected by the ageing process than by hearing sensitivity impairments. Further examination of Fig.4 shows that the SRT_HP_ was generally worse than the SRT_LP_. This effect was not influenced by the presence of background noise, suggesting that German speech intelligibility relies more on speech frequency information below 1.5 kHz.

In the LP condition, where hearing sensitivity was comparable and within the normal range for both the oNH and oHI groups, there were no significant differences in SRT_LP_. This supports the view that age-related differences were not the driving factor in explaining these group differences. In the HP condition, both the oNH and oHI groups had poorer SRT_HP_ scores than the yNH_control_ group, indicating a predominant age effect. Whereas the SiN condition was similarly reduced in oNH and oHI listeners, the significantly worse performance of the oHI group compared to the oNH group in the SiQ condition underscores the additional impact of reduced hearing sensitivity at frequencies above 1.5 kHz for SiQ, but not SiN, processing.

### Speech reception thresholds: individual differences

Figures 5 and 6 depict the relationship between RAM (top) or SAM (bottom) EFR amplitudes and SRT_SiQ_ or SRT_SiN_, respectively. Overall, the SRT related most strongly to the RAM, not SAM, EFR amplitude. After correcting for multiple comparisons (n=12, p=.0042), all SRT_SiQ_ conditions correlated significantly to the RAM EFR, while for the SRT_SiN_ conditions, only the BB and HP condition remained significant. This suggests that the RAM-EFR marker, which was more sensitive to detect individual CS differences than the SAM-EFR, was also more effective at predicting individual differences in speech recognition.

**Figure 5:**
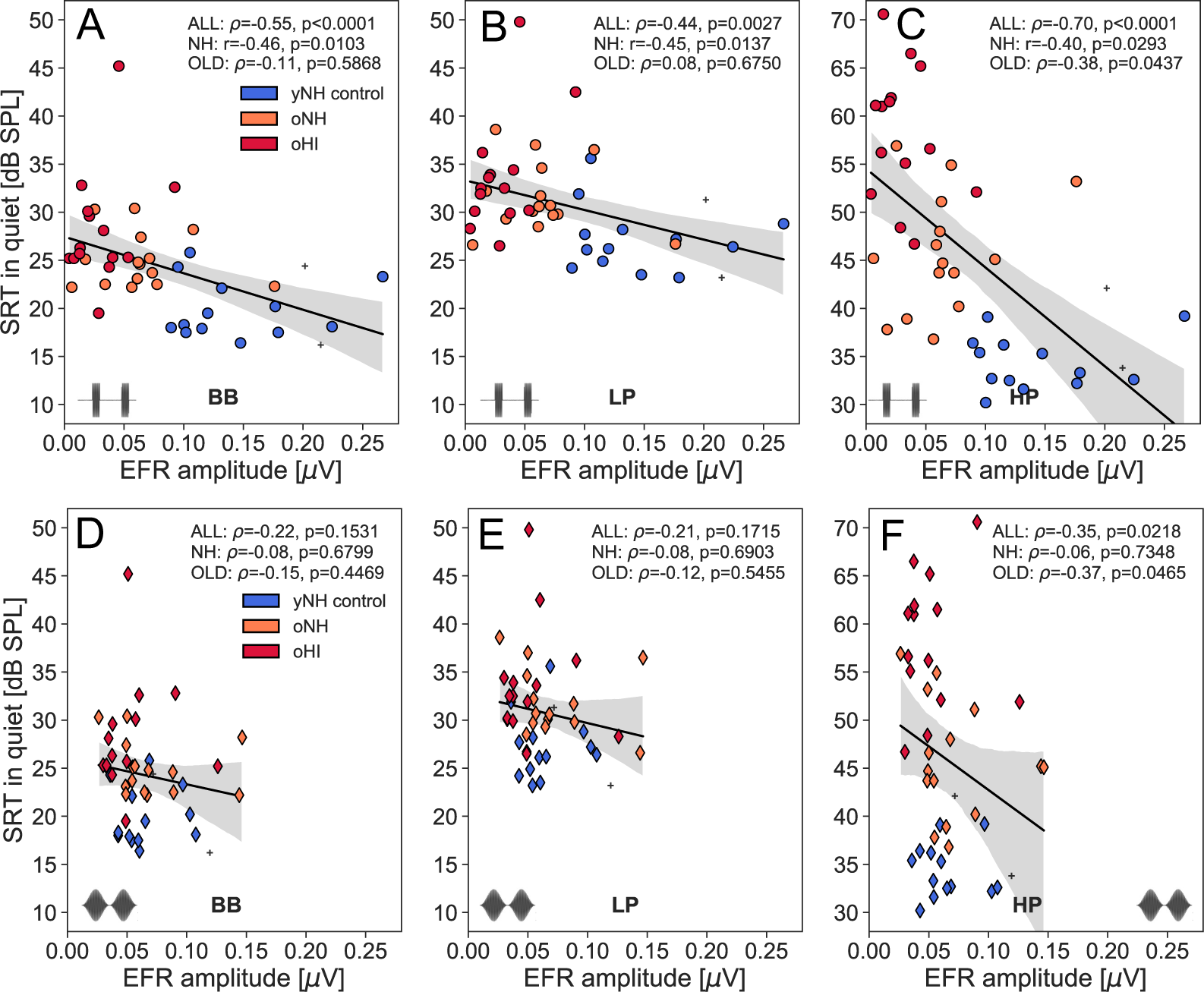
Regression plots between SRT_SiQ_ and the EFR amplitudes for RAM (top) and SAM stimuli (bottom). Analyses are performed for the BB **(A,D),** LP (**B,E**) and HP filtered conditions (**C,F**). Subjects belonging to the yNH_control_, oNH, oHI groups are colorcoded and the two yNH subjects who did not meet the TH_DP_ criterion to be included in the yNH_control_ group are marked with crosses. Correlation statistics (*ρ* or r) are indicated on the each panel and are performed across the entire cohort (ALL), or subgroups of OLD (oNH+oHI) or NH (yNH+oNH) subjects.

**Figure 6:**
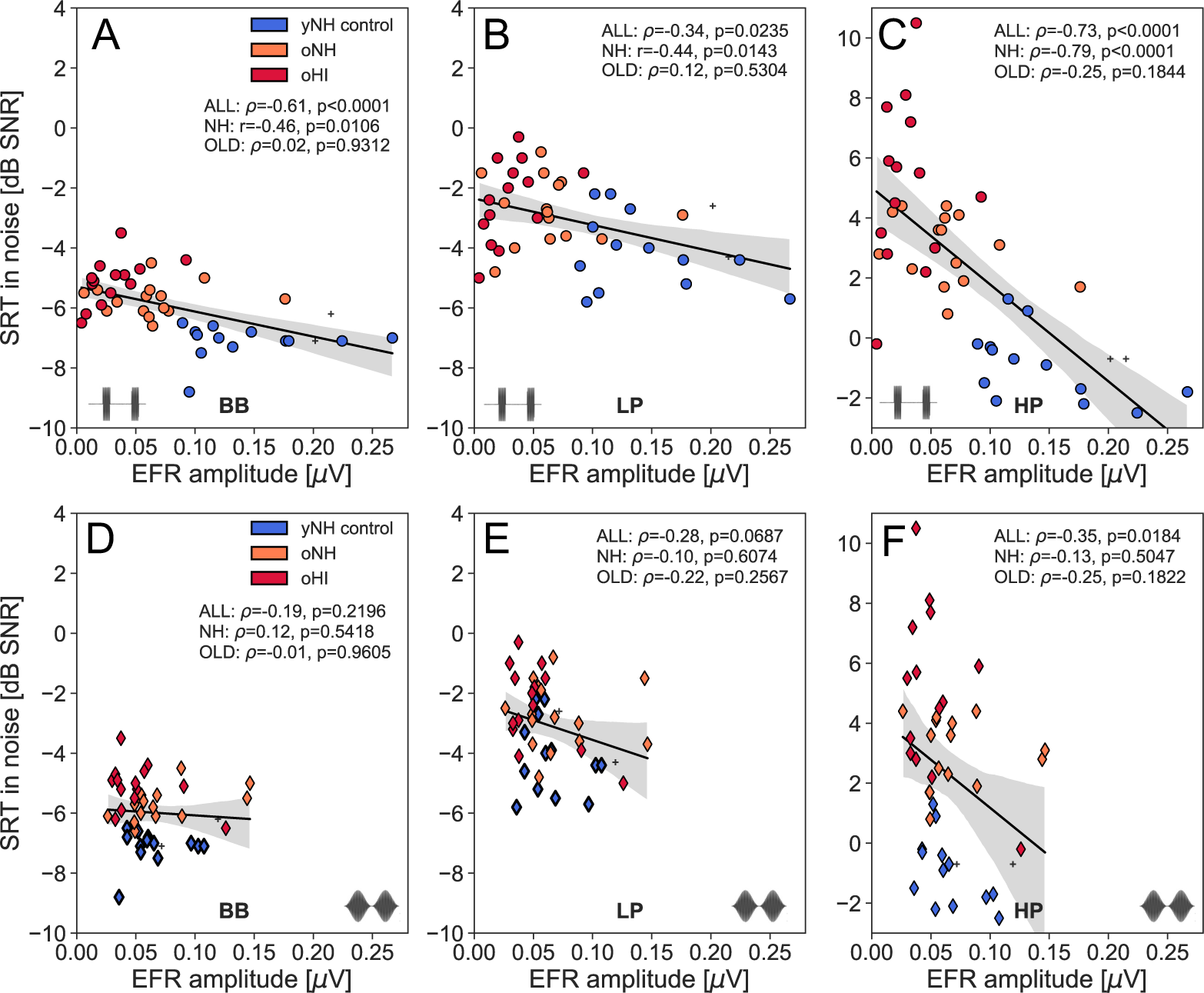
Regression plots between SRT_SiN_ and the EFR amplitudes for RAM (top) and SAM stimuli (bottom). Analyses are performed for the BB (**A,D**), LP (**B,E**) and HP filtered conditions (**C,F**). Subjects belonging to the yNH_control_, oNH, oHI groups are color-coded and the two yNH subjects who did not meet the TH_DP_ criterion to be included in the yNH_control_ group are marked with crosses. Correlation statistics (*ρ* or r) are indicated on the each panel and are performed across the entire cohort (ALL), or subgroups of OLD (oNH+oHI) or NH (yNH+oNH) subjects.

Secondly, the EFR markers specifically targeted TENV coding mechanisms, given their 4-kHz carrier frequency, which led to a stronger prediction of the SRT_HP_ than the SRT_LP_.

Table 1 and the figure legends summarize the correlation statistics between SRT conditions and the RAM EFR for the entire cohort (ALL) as well as for the NH or OLD subgroups. When comparing correlations across subgroups of NH and OLD participants, it is evident that the RAM EFR consistently serves as a stronger predictor of SRT in the NH cohort compared to the OLD cohort. This finding suggests that age-related effects, which are the dominant factor in the NH group, exert a greater influence on SRT predictions than TH_A_ differences do in the OLD cohort. Notably, participants in the OLD group may already exhibit some degree of age-related CS, which could mask the relationship between RAM EFR and SRT.

**Table 1:**

|  | SRT<sub>BB</sub> | SRT<sub>LP</sub> | SRT<sub>HP</sub> |
| --- | --- | --- | --- |
| SiQ | p<.001 | p=.003 | p<.001 |
| | $\rho = -0.55$ | $\rho = -0.44$ | $\rho = -0.70$ |
| SiN | p<.001 | p=.024 | p<.001 |
| | $\rho = -0.61$ | $\rho = -0.34$ | $\rho = -0.73$ |

Correlation statistics between the RAM EFR marker and SRT<sub>SiQ</sub> and SRT<sub>SiN</sub> across the entire cohort (n=44) for different filtering conditions (BB, LP, HP).

Of particular interest is the NH group, who are not currently classified as clinically hearing-impaired based on their normal audiograms, yet can demonstrate both reduced RAM-EFR amplitudes and impaired SRTs. These findings highlight the importance of conducting more detailed assessments of these individuals beyond standard audiograms, especially when they report difficulties with hearing.

However, to support a hypothesis in which RAM EFR and SRTs are simultaneously influenced by an underlying CS cause, it is essential to rule out other factors that could have affected this relationship. Potential confounding factors include aspects that impact speech recognition (such as cognition) but do not affect the RAM EFR, or a predominance of other interrelated SNHL pathologies (such as OHC damage) that could drive both measures in this relationship.

### The roles of hearing sensitivity and age in predicting the SRT

To factor out the mediating effect of hearing sensitivity on the observed relationships between the RAM EFR and SRT, we corrected for TH_A_ by considering the residuals of a linear regression model between TH_A_ and SRT. Figure 7 shows the relation between the SRT residuals and the RAM EFR after correcting for the mean TH_A_ across the frequencies in the 0.125-8kHz, 0.125-1.5 kHz, and 1.5-8kHz intervals, for the BB, LP, HP conditions, respectively. After correcting for hearing sensitivity, the SRT_SiN-HP_ residuals remained significantly correlated for the NH subgroup (r(29)=-0.6, p*<*.001), and approached significance for the cohort (*ρ*(44)=-0.28, p=.07). We repeated this analysis using TH_DP,4kHz_ as the correction factor, and found that the significant relationship between RAM EFR and SRT_SiN-HP_ decreased from *ρ*(44)= -0.73 (p*<*.001) to *ρ*(44)= -0.34, but remained significant (p =.02). Taken together, these analyses show that the individual SRT_SiN-HP_ differences cannot solely be explained by hearing sensitivity differences. None of the SRT_BB_ or SRT_LP_ residuals maintained a relationship to the RAM EFR, after correcting for hearing sensitivity. This is not surprising as the RAM EFR marker of CS reflects supra-threshold TENV processing above the phase-locking limit, and the SRT_BB_ values were very similar to the SRT_LP_ results which rely on low-frequency hearing mechanisms.

**Figure 7:**
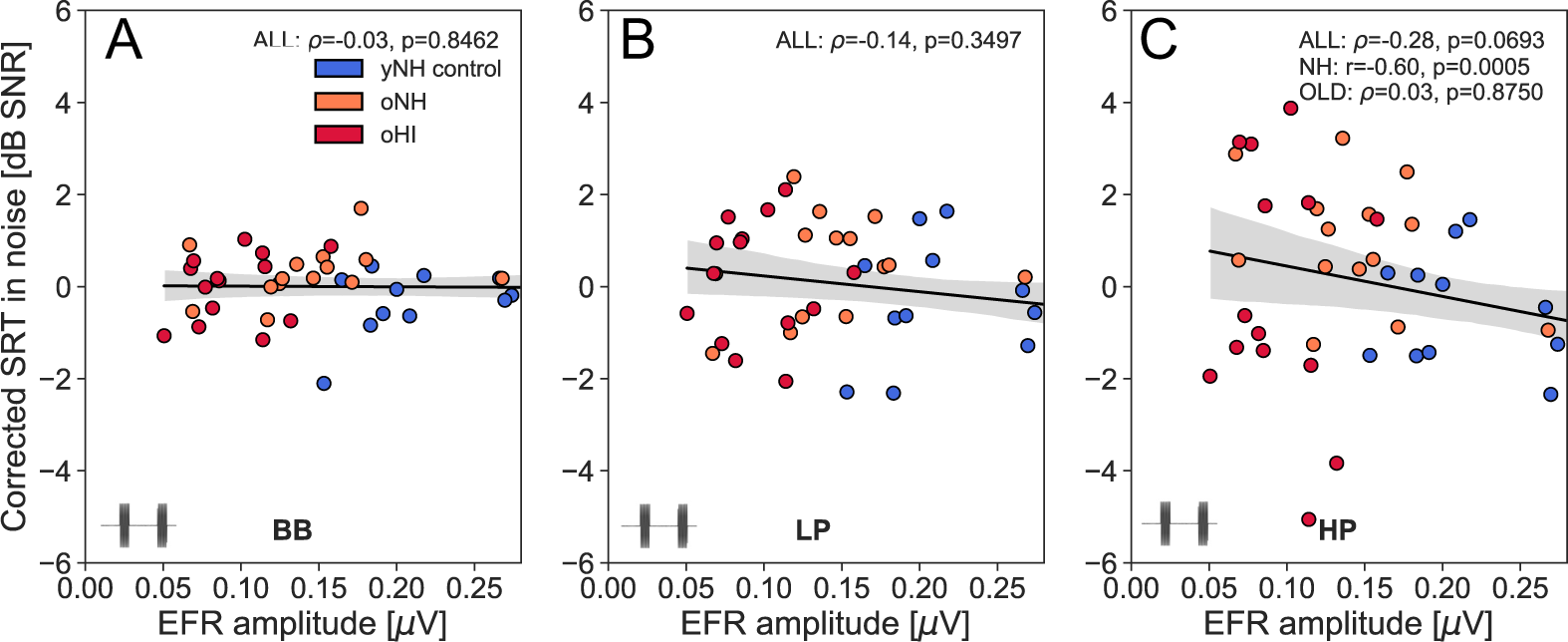
Residuals of a linear regression model between the SRT and TH*_A_* for the different SRT conditions: BB, LP, HP. The applied TH*_A_* for each of the conditions was adjusted to the frequency range of the speech material. The TH*_A_* correction applied refers to the mean TH*_A_* across the 0.125-8 kHz (BB), 0.125-1.5 kHz (LP), and 1.5 - 8kHz (HP) intervals, respectively. Indicated correlation statistics refer to the *ρ* or *r* calculated between the TH*_A_*-corrected SRTs and the RAM EFR, and were either calculated across the entire cohort (ALL) or subgroups of NH (yNH+oNH) or OLD (oNH + oHI) subjects.

Another confounding factor, namely age, could have affected the SRT and RAM EFR differently, hence we first considered its independent contribution to the RAM EFR. While a significant age effect was evident in the RAM EFR for the pooled data (*ρ*(44) = -0.66, p*<*.001). This effect was absent for the subgroup of older participants, who were age-matched within an 61-68 age bracket. Simultaneously, an age effect was also observed for the yNH_control_ subgroup (*ρ*(13) = -0.57, p = .041). These findings suggest that while a general age-related trend for CS exists, individuals within the same age decade can still exhibit varying degrees of CS.

Secondly, we investigated the independent contributions of age and TH to the SRT_HP_ as part of a linear regression analysis performed in Table 2. Age was a strong predictor of the SRT_HP_ in quiet and noise conditions (Adjusted R^2^ of 0.58 and 0.54, resp.), but is not an independent predictor as such. A colinearity analysis with age and other predictor values for the SRT (i.e., RAM EFR, TH_A_, TH_DP_) showed variance inflation factors above 1.5, indicating mild colinearity between the predictor variables. A Durbin-Watson test further revealed that linear regression models with age and TH_DP_, or age and RAM EFR had significantly correlated residuals (p*<*.05). Since age was neither independent of the RAM-EFR nor TH variables, including it in the multiple regression models would consistently dominate the results. Moreover, because age cannot be considered a marker of peripheral hearing damage, it was excluded from further analysis.

**Table 2:**
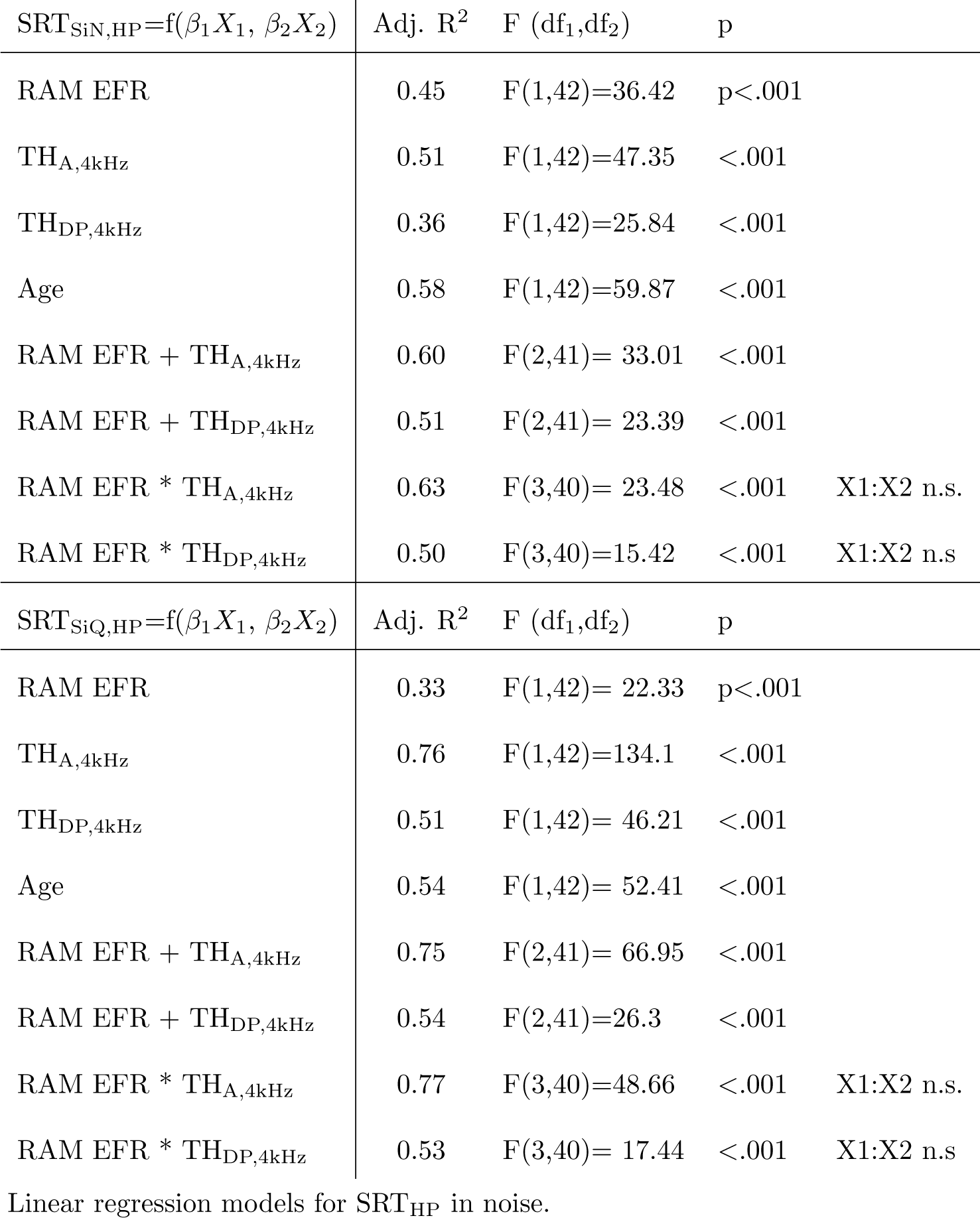

| $SRT_{SiN,HP}=f(\beta_1 X_1, \beta_2 X_2)$ | Adj. $R^2$ | F (df<sub>1</sub>,df<sub>2</sub>) | p | |
| --- | --- | --- | --- | --- |
| RAM EFR | 0.45 | F(1,42)=36.42 | p<.001 |  |
| TH<sub>A,4kHz</sub> | 0.51 | F(1,42)=47.35 | <.001 |  |
| TH<sub>DP,4kHz</sub> | 0.36 | F(1,42)=25.84 | <.001 |  |
| Age | 0.58 | F(1,42)=59.87 | <.001 |  |
| RAM EFR + TH<sub>A,4kHz</sub> | 0.60 | F(2,41)= 33.01 | <.001 |  |
| RAM EFR + TH<sub>DP,4kHz</sub> | 0.51 | F(2,41)= 23.39 | <.001 |  |
| RAM EFR * TH<sub>A,4kHz</sub> | 0.63 | F(3,40)= 23.48 | <.001 | X1:X2 n.s. |
| RAM EFR * TH<sub>DP,4kHz</sub> | 0.50 | F(3,40)=15.42 | <.001 | X1:X2 n.s |
| $SRT_{SiQ,HP}=f(\beta_1 X_1, \beta_2 X_2)$ | Adj. $R^2$ | F (df<sub>1</sub>,df<sub>2</sub>) | p | |
| RAM EFR | 0.33 | F(1,42)= 22.33 | p<.001 |  |
| TH<sub>A,4kHz</sub> | 0.76 | F(1,42)=134.1 | <.001 |  |
| TH<sub>DP,4kHz</sub> | 0.51 | F(1,42)= 46.21 | <.001 |  |
| Age | 0.54 | F(1,42)= 52.41 | <.001 |  |
| RAM EFR + TH<sub>A,4kHz</sub> | 0.75 | F(2,41)= 66.95 | <.001 |  |
| RAM EFR + TH<sub>DP,4kHz</sub> | 0.54 | F(2,41)=26.3 | <.001 |  |
| RAM EFR * TH<sub>A,4kHz</sub> | 0.77 | F(3,40)=48.66 | <.001 | X1:X2 n.s. |
| RAM EFR * TH<sub>DP,4kHz</sub> | 0.53 | F(3,40)= 17.44 | <.001 | X1:X2 n.s |

Linear regression models for $SRT_{HP}$ in noise.

Tables 2 and 3 report a multiple regression and commonality analyses which considered the entire cohort (n=44) and variables: RAM EFR, TH_A_, TH_DP_, or age using the following equation for the dependent variable (SRT_HP_):

**Table 3:**
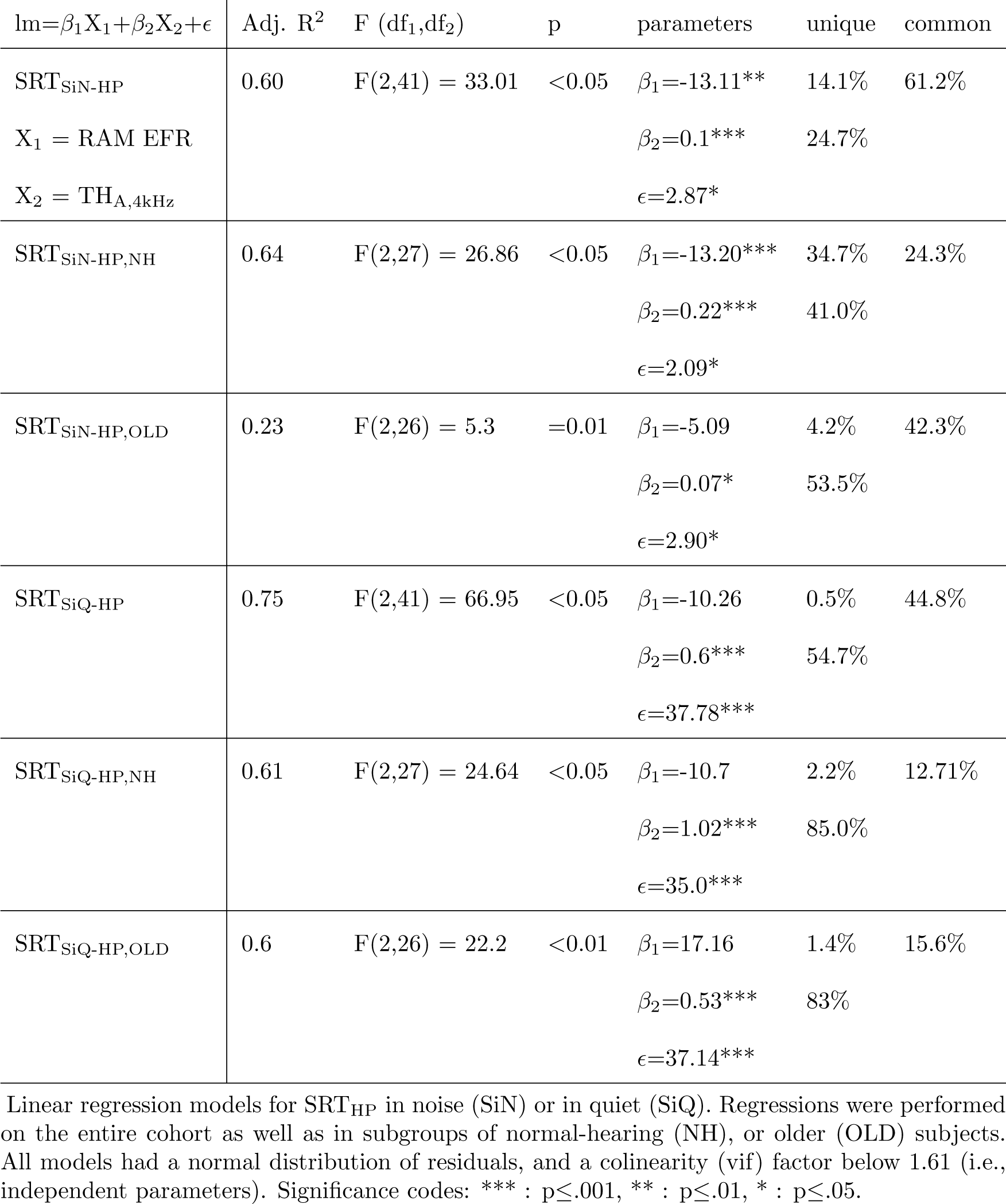

| $lm=\beta_1X_1+\beta_2X_2+\epsilon$ | Adj. $R^2$ | F (df<sub>1</sub>,df<sub>2</sub>) | p | parameters | unique | common |
| --- | --- | --- | --- | --- | --- | --- |
| SRT<sub>SiN-HP</sub> | 0.60 | F(2,41) = 33.01 | <0.05 | $\beta_1=-13.11^{**}$ | 14.1% | 61.2% |
| $X_1 = \text{RAM EFR}$ | | | | $\beta_2=0.1^{***}$ | 24.7% | |
| $X_2 = \text{TH}_{A,4\text{kHz}}$ | | | | $\epsilon=2.87^*$ | | |
| SRT<sub>SiN-HP,NH</sub> | 0.64 | F(2,27) = 26.86 | <0.05 | $\beta_1=-13.20^{***}$ | 34.7% | 24.3% |
| | | | | $\beta_2=0.22^{***}$ | 41.0% | |
| | | | | $\epsilon=2.09^*$ | | |
| SRT<sub>SiN-HP,OLD</sub> | 0.23 | F(2,26) = 5.3 | =0.01 | $\beta_1=-5.09$ | 4.2% | 42.3% |
| | | | | $\beta_2=0.07^*$ | 53.5% | |
| | | | | $\epsilon=2.90^*$ | | |
| SRT<sub>SiQ-HP</sub> | 0.75 | F(2,41) = 66.95 | <0.05 | $\beta_1=-10.26$ | 0.5% | 44.8% |
| | | | | $\beta_2=0.6^{***}$ | 54.7% | |
| | | | | $\epsilon=37.78^{***}$ | | |
| SRT<sub>SiQ-HP,NH</sub> | 0.61 | F(2,27) = 24.64 | <0.05 | $\beta_1=-10.7$ | 2.2% | 12.71% |
| | | | | $\beta_2=1.02^{***}$ | 85.0% | |
| | | | | $\epsilon=35.0^{***}$ | | |
| SRT<sub>SiQ-HP,OLD</sub> | 0.6 | F(2,26) = 22.2 | <0.01 | $\beta_1=17.16$ | 1.4% | 15.6% |
| | | | | $\beta_2=0.53^{***}$ | 83% | |
| | | | | $\epsilon=37.14^{***}$ | | |

Linear regression models for SRT<sub>HP</sub> in noise (SiN) or in quiet (SiQ). Regressions were performed on the entire cohort as well as in subgroups of normal-hearing (NH), or older (OLD) subjects. All models had a normal distribution of residuals, and a colinearity (vif) factor below 1.61 (i.e., independent parameters). Significance codes: \*\*\* : $p \leq .001$ , \*\* : $p \leq .01$ , \* : $p \leq .05$ .

**Table 4:**

| SAM EFR level<br>[dB SPL] | t(d.f.) | p |
| --- | --- | --- |
| 60 | t(12) = 1.49 | p = .16 |
| 65 | t(13) = 1.20 | p = .29 |
| 70 | t(14) = 1.61 | p = .13 |
| 75 | t(15) = 0.67 | p = .50 |
| 80 | t(17) = 1.24 | p = .23 |

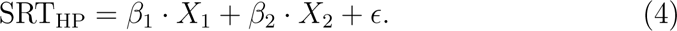

Several linear regression models of SRT_SiN,HP_ and SRT_SiQ,HP_ were compared: single regression models which only considered the X_1_ factor, and multiple regression models with and without interaction terms. Best-fitting single regression models were those with age or TH_A_ as the predictor variable. Regression models which included TH_A_ or TH_DP_ and the RAM EFR improved the model fit for the SRT_SiN,HP_, but not for SRT_SiQ,HP_ condition, and adding an interaction term did not further improve the models.

In general, hearing sensitivity parameters were stronger predictors in the SRT_SiQ,HP_ than SRT_SiN,HP_ models. This underscores a strong association between SRT_SiQ_ and hearing sensitivity (Festen & Plomp, 1983; Papakonstantinou et al., 2011). In contrast, it is well known that the SRT_SiN_ is not well predicted by hearing sensitivity alone (Plomp, 1986; Fitzgerald et al., 2024). The SRT_SiN-HP_ performance may thus be more influenced by suprathreshold TENV deficits linked to temporal coding impairments caused by CS (Lopez-Poveda & Barrios, 2013) or so-called supra-threshold distortion factors (Plomp, 1986).

To further examine the independent role of EFR RAM for speech recognition, we performed linear regression models and a commonality analysis for the dependent variables SRT_SiN,HP_ and SRT_SiN,HP_ in Table 3. All models had a normal distribution of residuals, and a colinearity (vif) factor below 1.61 (i.e., *<*5, thus independent variables). Homoscredasticity was not met for the SRT_SiN,HP_ models, and the Durbing-Watson test showed dependence in the SRT_SiN,HP_ and SRT_SiN,NH_ conditions. Linear models with RAM EFR (X_1_) or TH_A,4kHz_ (X_2_) were significant for all conditions, but explained less variance for the SRT_SiN-HP, OLD_ condition where only minor age-related CS differences were expected due the age-matching of subjects in the cohort. Even though there was a significant unique contribution of TH_A,4kHz_ in all models, there was also a unique and significant contribution of 14.1% and 34.7% for the RAM EFR in the SRT_SiN,HP_ and SRT_SiN-HP,NH_ models, respectively. Repeating this analysis, but with the TH_DP,4kHz_ variable of hearing sensitivity, yielded unique RAM EFR contributions of 30.37% and 82.59 % for the SRT_SiN-HP_ and SRT_SiN-HP,NH_ conditions, respectively. Especially given that the unique contribution of RAM EFR reduced drastically in the SRT_SiQ,HP_ models, adds weight to the conclusion that our proposed RAM EFR marker of CS is more effective at predicting the individual signal-to-noise ratio required for speech recognition in a fixed-level, stationary noise background.

## Discussion

We report strong predictive power of the RAM-EFR amplitude for SRTs, with better performance for SRT_HP_ compared to SRT_BB_ and SRT_LP_ conditions (Table 1, Figs.5, 6). Additionally, the unique contribution of the RAM EFR marker of CS to the SRT_HP_ models, which included both hearing sensitivity and the RAM-EFR marker, was greater for the SiN condition than for the SiQ condition (Table 3). This finding aligns with the prevailing hypothesis that CS has a more pronounced effect on supra-threshold SiN processing than on SiQ processing. Overall, our results suggest that the ANF population, as assessed using a fixed-level 70-dB-SPL, 4-kHz RAM stimulus, is a strong predictor of speech recognition at similar sound levels and within comparable cochlear frequency ranges.

### The quality of the adopted RAM-EFR marker of CS

To interpret the RAM-EFR as a pure marker of CS, the EFR marker would need to be fully independent of hearing sensitivity or OHC integrity. Our recordings from Budgerigars support the sensitivity of the EFR marker to CS, though they do not completely exclude the possibility that significant OHC damage may also influence its amplitude. Model simulations presented by Vasilkov et al. (2021) provide further insight, indicating that on-CF ANF responses to the RAM stimulus remain unaffected by OHC damage (see their Fig.1). Additional simulations of human EFR generators by Vasilkov et al. (2021); Van Der Biest et al. (2023) show that simulated OHC damage has only a minimal effect (5–10%) on the 4-kHz RAM EFR amplitude, while CS significantly impacts the response, reducing it by up to 81%. Thus, the RAM stimulus was designed to be minimally influenced by coexisting OHC damage, a conclusion supported by our human EFR recordings, which show a greater EFR amplitude difference between the yNH_control_ and the oNH group than between the oNH and oHI groups (see Fig.3D).

We argue that when the stimulus is presented at sufficiently high levels to drive ANFs into saturation, the impact of the effective stimulus level, or supra-threshold audibility, on the EFR response is minimal. Although this effect was not systematically examined in our study, we reference the findings of Encina-Llamas et al. (2021). In their study, EFRs were recorded in response to 98-Hz-modulated SAM tones at various pure-tone frequencies and levels in young normal-hearing listeners (mean age: 24 *±* 3.2 years) and older hearing-impaired listeners (mean age: 56.2 *±* 12.7 years). The hearing-impaired participants had hearing thresholds of *<*=20 dB HL below 4 kHz and between 20 and 45 dB HL for frequencies up to 8 kHz, which aligns closely with the audiometric profiles considered in our study.

By fitting two piecewise linear curves to the EFR magnitude growth curves between 20 and 80 dB SPL (expressed in dB per dB), Encina-Llamas et al. (2021) demonstrated that both NH and HI growth curves exhibited a compression breakpoint around 60 dB SPL (see Fig. 2E in their study). To examine this further, we replotted their original EFR data in *µ*V to calculate growth slopes for stimulus levels above 60 dB SPL, as shown in Fig.8. Our analysis revealed an EFR growth slope of 0.002 *µ*V per dB, which was similar for both NH and HI listeners. This indicates that EFR amplitudes increase by approximately 0.04 *µ*V between stimulus levels of 60 and 80 dB SPL, and that differences in hearing sensitivity did not affect this process.

**Figure 8:**
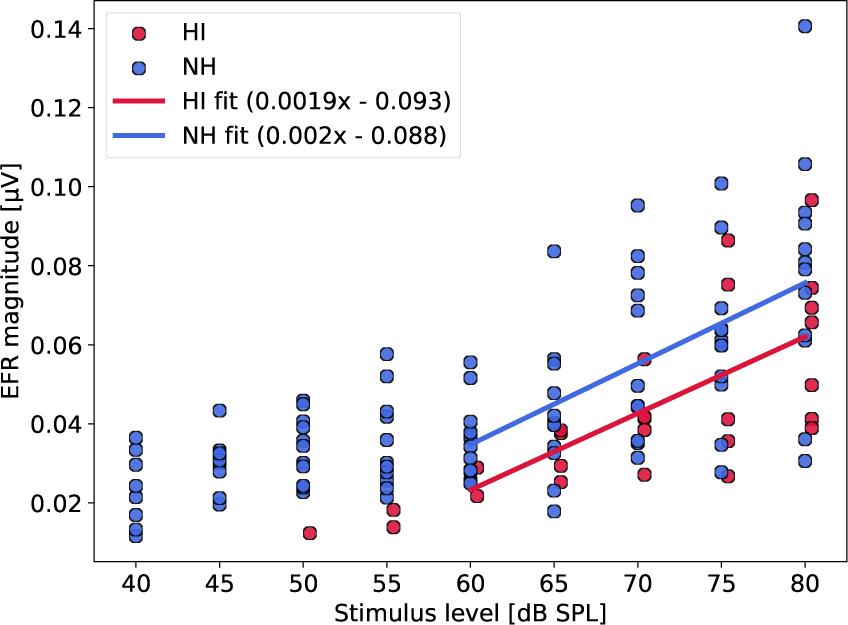
Reanalysis of the Encina-Llamas, et al. (2017) dataset reporting EFR magnitudes to a 98-Hz SAM tone of 4011 Hz. dB values were transformed into using a 10**(dB/20) transformation with 1 V as the reference. Data from 13 NH (24 *±* 3.2 y/o) and 7 HI (56.2 *±* 12.7 y/o) listeners are shown. Only EFR amplitudes that were significantly above the noise floor are shown, and a linear fit was made across data-points above the compression knee-point of 60 dB SPL. The NH cohort had audiogram thresholds below 15 dB HL for frequencies below 8 kHz and the HI cohort had dB HL thresholds *<*= 20 for frequencies below 4 kHz and between 20 and 45 dB HL for frequencies up to 8 kHz.

More importantly for our study, the dataset from Encina-Llamas et al. (2017) did not show significant group differences in EFR amplitude between NH and HI listeners at stimulation levels of 60 dB SPL or higher (see Table 4). Hearing sensitivity differences of up to 30 dB between NH and HI subjects for frequencies above 4 kHz did thus not significantly impact the supra-threshold EFR amplitude. This suggests that variations in audibility are unlikely to influence EFR amplitude in terms of supra-threshold TENV coding or CS when presented at a fixed, supra-threshold level above the EFR growth curve’s knee-point.

Taken together, the model simulations (Vasilkov et al., 2021; Van Der Biest et al., 2023) and experimental findings from both Budgerigar and human studies support the conclusion that our EFR marker is sensitive to CS and is largely independent of hearing sensitivity differences when assessed at 70 dB SPL. Our human results demonstrated a clear age-related decline in EFR amplitudes, even in the absence of OHC damage. These findings align with animal research linking deficits in temporal coding at the earliest neural stages of the auditory pathway to progressive or noise-induced CS (Parthasarathy et al., 2014; Fernandez et al., 2015; Shaheen et al., 2015; Parthasarathy & Kujawa, 2018).

### Context with prior studies

This study addressed and minimized several challenges that have complicated previous human research into the causal relationship between speech-in-noise intelligibility deficits and EFR markers of synaptopathy. One key issue in earlier studies has been the low sensitivity of the traditionally used SAM-based EFR metrics (Grose et al., 2017; Guest et al., 2018). In contrast, we demonstrate that the RAM-EFR is consistently larger in the same individuals and is more strongly affected by KA-induced CS than the conventional SAM-EFR (Fig.3). This increased sensitivity makes the RAM-EFR more effective than the SAM-EFR for detecting individual differences in both CS and SRTs (Figs.5 and 6).

Additionally, prior model simulations have shown that the 4-kHz SAM EFR is more influenced by OHC damage compared to the RAM EFR, highlighting the superior specificity of the RAM EFR for detecting CS (Vasilkov et al., 2021; Van Der Biest et al., 2023). By integrating these simulations with Budgerigar and human EFR recordings, we provided evidence that the two variables considered in the regression models—hearing sensitivity and the RAM-EFR marker of CS—can be treated as independent factors of SNHL.

Furthermore, by better aligning the frequency content of the EFR and speech stimuli to target auditory TENV mechanisms and their deficits, the models we presented are better suited than previous studies to isolate the independent contribution of CS to speech intelligibility in noise.

### Confounding factors and Study limitations

Since synaptopathy often precedes permanent OHC damage during the aging process (Sergeyenko et al., 2013; Fernandez et al., 2015; Parthasarathy & Kujawa, 2018) or following noise exposure (Kujawa & Liberman, 2009; Furman et al., 2013), markers of OHC damage may inadvertently serve as predictors of CS. Although these two pathologies are distinct in their origins and have different effects on auditory processing, they share common risk factors, such as age and noise exposure, and both are associated with reduced EFR amplitudes (Dimitrijevic et al., 2016; Parthasarathy & Kujawa, 2018; Garrett & Verhulst, 2019). This complexity underscores the importance of ensuring that a marker for CS is independent of OHC damage markers (e.g., TH_DP_ or TH_A_) when assessing the specific role of CS in sound perception. While we provide strong evidence that the RAM-EFR marker primarily captures CS-related information, its effectiveness may still correlate with auditory threshold measures if individuals with reduced hearing sensitivity also experience CS as an early sign of SNHL.

Our multiple regression analysis revealed that SRTs are influenced by both CS and behavioral and DPOAE markers of hearing sensitivity (Table 3). In the oHI subgroup, the lack of a clear relationship between SRT_SiN_ and the RAM EFR may be explained by the presence of OHC damage (Fig.6C). While CS contributed to SRT deficits in all listeners within this subgroup, OHC damage likely exacerbated the age-related decline in SRT without being reflected in the EFR marker. Additionally, the group-level analysis (Fig.4B) underscores the compounded impact of OHC damage on speech intelligibility in noise, as evidenced by significantly worse SRT_SiN-HP_ performance in the oHI group compared to the yNH_control_ group. Lastly, our results show that the oNH group can perform as poorly on SiN processing as the oHI listeners, and this has important consequences for clinical practice. While the oNH group is currently not considered to have a hearing problem based on their audiogram, they do experience speech processing deficits that should be quantified in clinical practice using either a SiN test, or an EFR marker of CS which predicts their SRT scores.

Dissociating general age effects from age-related CS remains challenging in human studies. Previous research involving older participants or those with significant noise-exposure histories has often reported electrophysiological evidence of CS or reduced temporal coding fidelity in the studied populations (e.g. Anderson et al., 2011, 2012; Konrad-Martin et al., 2012; Clinard & Tremblay, 2013; Schoof & Rosen, 2016; Bramhall et al., 2017; Valderrama et al., 2018; Bramhall et al., 2021). Within this context, our study confirms that age is inherently linked to the development of SNHL. However, in our age-restricted samples (OLD or yNH_control_ subgroups), age proved to be a poor predictor of the degree of CS. More broadly, age is associated with both OHC damage (Lin et al., 2011; ISO, 2017) and CS (Schmiedt et al., 1996; Makary et al., 2011; Konrad-Martin et al., 2012; Möhrle et al., 2016; Parthasarathy & Kujawa, 2018). This dual association likely contributes to the observed strong relationship between age and RAM-EFR when pooling data across groups (*ρ*(44) = -0.66, p *<*.001). Because age correlates with all facets of SNHL and its biomarkers, incorporating it into statistical models complicates the interpretation of the unique contributions of each pathology.

Next, we address several aspects related to our study design. Current limitations in human experimentation prevent us from directly establishing a causal relationship between CS and speech intelligibility in live humans, necessitating reliance on indirect evidence to support our conclusions. To this end, we incorporated model simulations that examined the impact of various SNHL pathologies on the cochlear and neuronal generators of the EFR (Vasilkov et al., 2021; Van Der Biest et al., 2023). Additionally, we employed an animal model to demonstrate the sensitivity of our EFR markers to CS. The convergence of findings from model predictions and experimental results in several studies (e.g. Verhulst et al., 2018; Encina-Llamas et al., 2019; Keshishzadeh et al., 2020; Vasilkov et al., 2021; Buran et al., 2022) supports the validity of these approaches for the purposes of our study. However, it is important to acknowledge that current state-of-the-art models, while advanced, may not fully capture all cochlear and neural mechanisms underlying EFR generation. Moreover, species differences could confound direct comparisons between Budgerigar and human EFRs. Nevertheless, neural recordings from the Budgerigar inferior colliculus indicate that their auditory processing strategies are comparable to those of the mammalian midbrain (Henry et al., 2017), supporting the relevance of these findings to human research.

In our approach to isolate the contribution of CS to speech intelligibility deficits within the broader context of SNHL, we conducted a group analysis. This approach aimed to amplify the effects between the yNH_control_ group and older groups expected to exhibit age-related CS. However, pooling data from groups that differ in multiple factors can introduce additional between-group explanatory variables that were not explicitly controlled. For instance, cognitive factors such as memory and attention, which are known to decline with age and are associated with speech-in-noise comprehension (Humes et al., 2010; Humes, 2013; Yeend et al., 2017), likely account for some of the unexplained variance in the multiple regression models (Table 3). This is particularly relevant as the analyzed groups had substantial age differences. However, the matrix test was designed to mitigate memory effects by randomly generating word sequences and minimizing cognitive load through the immediate recall of only five words at a time. Additionally, EFR recordings in response to high-modulation-frequency stimuli are considered largely free from top-down attention (e.g. Varghese et al., 2015) or memory influences. The robust correlations observed between RAM-EFR amplitudes and SRT_SiN-HP_ scores suggest that cognitive factors are not the primary driving force behind the findings, but instead interact with peripheral auditory encoding deficits (Johannesen et al., 2016). We contend that, for the purposes of this study, pooling data from carefully defined homogeneous groups is both valid and necessary to disentangle the relative contributions of OHC damage and CS to speech intelligibility deficits. Future studies which consider an age-gradient across a larger cohort of study participants may further shed light on the dynamics between age, CS and speech intelligibility.

We propose the following considerations for future study designs. Recent research on ANF coding (Henry et al., 2016; Encina-Llamas et al., 2019; Vasilkov & Verhulst, 2019) has demonstrated that for supra-threshold stimulation, TENV information can be encoded via the tails of high-spontaneousrate ANF tuning curves in basal cochlear regions. This suggests that the neural generators of the 4-kHz EFR may encompass a broader frequency range than expected from the stimulus’ narrow-frequency profile and its basilar membrane excitation. Such a phenomenon could impact the intended alignment between the cochlear frequency regions activated by the EFR marker and those involved in processing speech stimuli, particularly under the assumption that both are governed by similar TENV mechanisms. While our study design improved this alignment by transitioning from the broadband (BB) condition to the high-pass (HP) condition, additional refinements could further enhance the interpretation of how reduced TENV coding contributes to speech intelligibility deficits. For instance, introducing high-frequency masking noise to both the speech and EFR stimuli could better isolate the frequency regions of interest and improve alignment.

### Conclusion

We conclude that age-related synaptopathy is a significant hearing health concern, as both experimental and theoretical evidence demonstrate its substantial impact on auditory TENV coding and speech intelligibility, especially in noise. Sensitive diagnostic tools are essential for understanding the role of synaptopathy in impaired sound perception. The RAM-EFR marker, which proved to be robust, widely applicable, and selective for CS, was instrumental in reaching these conclusions. Given the independent contribution of synaptopathy to age-related speech perception deficits, particularly in noisy environments, future therapeutic interventions can be developed to address synaptopathy and mitigate its functional consequences.

## Abbreviations

ABR: auditory brainstem response
AEP: auditory evoked potential
AN(F): auditory nerve (fiber)
BB: broadband
CAP: compound action potential
CF: characteristic frequency
DPOAE: distortion product otoacoustic emission
EEG: electroencephalography
EFR: envelope following response
HI: hearing impaired
HP: high-pass filtered
IHC: inner hair cell
KA: Kainic Acid
LP: low-pass filtered
NH: normal hearing
OHC: outer hair cell
peSPL: peak-equivalent sound pressure level
PT: pure-tone
SAM: sinusoidally amplitude-modulated
SiN: speech in noise
SiQ: speech in quiet
SNHL: sensorineural hearing loss
SNR: signal-to-noise ratio
SRT: speech reception threshold
SPL: sound pressure level
RAM: rectangularly amplitude-modulated
TENV: temporal envelope
TFS: temporal fine structure
TH_A_: audiometric hearing threshold
TH_DP_: distortion-product otoacoustic emission threshold

## Acknowledgments

This work was supported by the DFG Cluster of Excellence EXC 1077/1 “Hearing4all” (MG, MM, SV), the European Research Council (ERC) under the Horizon 2020 Research and Innovation Programme (grant agreement No 678120 RobSpear; VV, SV), National Institutes of Health grant R01 DC017519 (KH) and a National Institutes of Health Predoctoral National Research Service Award Fellowship (TL1 TR002000) administered by the University of Rochester Clinical and Translational Science Institute (JW). The authors would like to thank the study participants as well as the Hörzentrum Oldenburg for helping with participant recruitment. Lastly, we thank Sarineh Keshishzadeh for help with the analysis scripts and data storage and labelling throughout the project and Attila Fŕater for help with the reanalysis of the Encina-llamas data.

## Author Contributions

MG: Conceptualization, Methodology, Software, Validation, Formal analysis, Investigation, Data Curation, Writing: Original Draft, Visualization; VV: Methodology, Software, Investigation; MM: Methodology, Software, Investigation; PDV: Statistics, Writing: Review & Editing; JW: Methodology, Software, Investigation; KH: Methodology, Software, Investigation; SV: Conceptualization, Methodology, Investigation, Resources, Writing: Original Draft, Writing: Review & Editing, Supervision, Project administration, Funding acquisition.

